# From sequences to schemas: low-rank recurrent dynamics underlie abstract relational representations

**DOI:** 10.64898/2026.04.10.717706

**Authors:** Vezha Boboeva, Alberto Pezzotta, George Dimitriadis, Athena Akrami

## Abstract

A hallmark of intelligent behavior is the ability to extract abstract relational structure from temporal sequences — recognizing, for instance, that `aab, ccd`, and `eef` all follow the same underlying pattern, regardless of the specific elements involved. This capacity, observed across species and sensory modalities, is thought to underlie the formation of cognitive schemas: compressed internal models that support rapid generalization to novel experiences. Yet the neural circuit mechanisms by which such abstract, identity-independent representations emerge from sequential experience remain largely unknown. Here, we investigate this question using Recurrent Neural Networks (RNNs) as mechanistic models of neural circuits, trained to classify sequences based on their latent algebraic patterns (e.g., `aab, aad → AAB; aba, aca → ABA`) without supervision on intermediate transitions. We demonstrate that RNNs spontaneously learn low-dimensional representations that mirror the hierarchical generative structure of the sequences, and that this abstraction is mechanistically supported by the emergence of low-rank recurrent connectivity. The leading singular component of the recurrent weight matrix integrates relational transition information — whether consecutive tokens are the same or different — across time, driving the formation of a structured, tree-like geometry in the population state space. Through singular vector ablation, we establish a causal role for this component: removing it selectively erases memory for earlier transitions while leaving local, single-step sensitivity intact. Finally, while RNNs trained on next-token prediction do not spontaneously acquire these abstract representations, transferring the low-rank scaffold learned from classification significantly accelerates learning and improves generalization — an effect specific to the abstract structure of the scaffold rather than generic statistical pretraining. These findings offer a computational account of how task demands shape recurrent connectivity to support temporal abstraction, with direct implications for understanding schema formation in biological brains.

## Introduction

### Schemas, abstraction, and the brain

A defining capacity of biological intelligence is the ability to form *schemas*: structured internal representations that capture the relational organization of experience independently of its specific sensory content [1, 2]. The concept of the schema, introduced by Bartlett, refers to organized knowledge structures that shape how new experiences are encoded, interpreted, and reconstructed from memory [3]. Rather than storing each experience as an isolated trace, neural circuits appear to extract relational structure shared across experiences and use it as a scaffold for encoding, retrieval, and generalization. Animals can assimilate new experiences into pre-existing schemas far faster than naive learning would permit, and this process depends critically on the hippocampus [4–6], medial prefrontal cortex [7, 8], and their interaction [9–11]. More broadly, the hippocampus has been implicated in building and maintaining relational, context-general representations [5, 6], whereas prefrontal cortex is thought to support the flexible deployment of abstract rules across contexts [7, 12].

Despite this progress, a central mechanistic question remains unresolved: *how do neural circuits construct abstract relational representations from sequential experience?* What circuit-level properties allow a population of neurons to represent not the identity of individual stimuli, but the pattern of relationships among them? And what determines whether such representations generalize to novel instantiations of familiar structure? Addressing these questions requires linking the cognitive notion of schema formation to the dynamics and connectivity of neural circuits.

### Statistical learning of sequential structure across species

A useful entry point into this problem is *statistical learning*: the spontaneous, often implicit acquisition of regularities embedded in sequential input, without explicit instruction or reward [13]. A particularly revealing form of such regularity is sensitivity to algebraic temporal patterns such as `AAB` or `ABA`, which specify identity relations among successive elements independently of their sensory content [14, 1]. A sequence of tones following an `ABA` pattern instantiates the same abstract rule as a sequence of shapes or words with the same relational structure, even though the sensory input differs completely. Sensitivity to such identity-independent patterns has been documented across a broad phylogenetic range, including humans across development [15, 16], non-human primates [15], rodents [17, 18], and multiple avian species [19–21]. This breadth suggests that algebraic pattern learning reflects a fundamental computational strategy rather than a language-specific capacity. At the same time, behavioral evidence alone does not reveal what kind of internal representation supports such generalization, nor how it is implemented by recurrent circuits [22].

### Population geometry and low-rank recurrence as candidate mechanisms

Recent work in systems neuroscience has begun to reveal how abstract information is organized in neural population activity. Across cortical and hippocampal circuits, collective activity during cognitive tasks often occupies a surprisingly low-dimensional subspace despite the high dimensionality of individual neuron responses [23, 24]. Crucially, the geometry of this subspace is not arbitrary: it reflects task structure. In prefrontal cortex, population trajectories during working memory and decision-making tasks trace out manifolds aligned with task-relevant variables [8, 25]. In hippocampus, population activity organizes into maps that encode relationships between experiences rather than isolated events [5, 26]. These observations suggest that abstraction may be expressed in the geometry of neural population states.

A natural theoretical framework for linking recurrent circuit structure to such geometry is the theory of low-rank recurrent networks [27, 28]. In this framework, the recurrent weight matrix is decomposed into a structured low-rank component, which confines activity to a task-relevant low-dimensional manifold, plus a random high-dimensional background. Each rank-one component defines a distinct computational channel, mapping a particular pattern of input selectivity to a corresponding recurrent output pattern. Because both Hebbian and gradient-based synaptic updates take the form of outer products, learning naturally builds low-rank structure on top of a random recurrent backbone. Low-rank recurrence is therefore an attractive candidate mechanism for the emergence of abstract representations from sequential experience.

### RNNs as models of neural circuit dynamics

Recurrent Neural Networks (RNNs) are well suited to modeling time-evolving processes such as working memory, context integration, and decision making [29], and are increasingly used to study how neural dynamics support behavior [30–32]. Their appeal as neural circuit models lies in the direct correspondence between recurrent synaptic interactions, external input, and nonlinear dynamics in biological circuits and in trained networks. RNNs trained on behavioral tasks reproduce a range of neural phenomena, including mixed selectivity, low-dimensional trajectories, and context-dependent dynamics, while remaining tractable enough to support mechanistic analysis [32]. In particular, the low-rank framework applies directly to trained RNNs, allowing one to link learned connectivity to the geometry of population activity.

### The present work

Here, we use RNNs to investigate the circuit mechanisms underlying the emergence of abstract relational representations from sequential experience. We ask two questions. First, what properties of recurrent connectivity allow a network to represent not the identity of individual tokens, but the abstract pattern of transitions among them? Second, what role does the task objective play in determining whether such representations emerge at all?

To address these questions, we generate algebraic sequences using a binary branching tree whose terminal nodes define abstract sequence classes (denoted by capital letters in this manuscript, e.g., `AAB` or `ABA`) and instantiate each class with concrete tokens drawn from a finite alphabet. We train RNNs to classify sequences according to their overall abstract pattern, receiving only the class label at the end of the sequence (e.g., `AAB` for `aac` and `bba`; `ABA` for `aca` and `ada`), without supervision on intermediate transitions. This design captures a key feature of natural statistical learning: the abstract rule is not signaled locally at each step, but must be inferred from the global structure of the sequence.

We show that trained networks generalize to novel token instantiations and develop low-dimensional hidden representations whose geometry mirrors the branching structure of the generative tree. We then identify a mechanistic basis for this geometry: recurrent connectivity becomes effectively low-rank, and its leading singular components support abstraction by integrating transition history across time, as revealed by rank-constrained training and singular-vector ablations. Finally, we test whether this learned structure is task-specific or reusable. Networks trained from scratch on next-token prediction do not develop the same hierarchical, low-rank organization. However, initializing prediction networks with classification-trained recurrent weights accelerates learning and improves generalization. Crucially, this benefit is specific to the learned abstract scaffold: control experiments using autoencoder pretraining fail to improve generalization, indicating that the transfer gain arises from structural schema rather than generic statistical pretraining. Together, these results provide a mechanistic account of how task demands shape low-rank recurrent dynamics to support relational abstraction and transfer in sequential computation, and they generate concrete predictions for the geometry of neural population activity in circuits engaged in schema learning.

## Previous Work

Understanding how neural systems represent and generalize sequential structure has long been a goal in both neuroscience and machine learning.

### Sequential memory and working memory in neural circuits

Early computational work focused on mechanisms for maintaining and recalling ordered information, especially in working memory, for instance by combining item and positional rank information [33]. More recently, RNNs have been shown to learn generalized positional structure [34], and related approaches use chunking or segmentation to extract interpretable units from high-dimensional neural dynamics, including structural schemas [35]. On the biological side, hippocampal place and time cells provide examples of circuits that encode the ordinal position of experience within a sequence [36], while prefrontal circuits maintain rule and context information across delays in a largely population-level, abstract format [25, 8].

### Cognitive maps, external memory, and sequence generalization

Another line of work augments RNNs with external memory or plasticity mechanisms, allowing sequences to be represented as trajectories over latent states or “cognitive maps” [6]. These approaches are biologically motivated, given strong evidence that the hippocampus constructs map-like representations that extend beyond physical space to encompass abstract relational structure [5, 26]. While powerful, such models often rely on local or Markovian transitions, and their ability to generalize to non-adjacent or higher-order dependencies remains unclear. Algebraic pattern learning poses a particularly strong version of this challenge because it requires representing not just the immediately preceding event, but the relational pattern linking the full sequence.

### Low-rank connectivity and low-dimensional neural dynamics

From a dynamical-systems perspective, low-rank recurrent connectivity is known to confine neural activity to task-specific low-dimensional manifolds [27, 28]. This framework has been used to analyze computations including working memory, decision-making, and context-dependent processing. Empirically, low-dimensional population structure is well established in cortical and hippocampal circuits [23], and there is growing evidence that task learning progressively sharpens this structure by reducing the effective dimensionality of activity and aligning it with task-relevant variables [24, 32]. Our work builds on this framework by showing how low-rank recurrent dynamics give rise specifically to *abstract relational* representations: not merely low-dimensional activity, but a population geometry that encodes transition structure independently of stimulus identity. In contrast to prior studies focused on positional order or explicit memory architectures, we show that standard RNNs can spontaneously learn algebraic relational patterns through low-rank dynamics alone, and we identify a causal role for the dominant singular components in integrating transition history to form hierarchical representations.

## 1 Results

Our approach is to use RNNs as tractable models of recurrent neural circuits, and to ask what circuit properties spontaneously emerge when such a network must solve a problem requiring abstract temporal generalization. Specifically, we are interested in three interrelated questions: (1) what is the geometry of the population activity that supports identity-independent sequence classification; (2) what property of the recurrent connectivity gives rise to that geometry; and (3) what role does the task objective play in determining whether abstract representations form at all.

Each of these models has a standard discrete-time RNN as a core component. The inputs to the RNN, denoted ***x*** ∈ ℝ^*α*^, are a one-hot encoding of the elements in the sequence over letters in the alphabet. The *µ*-th sequence, ***X***^*µ*^, with *µ* = 1 … *p*, is given by the elements 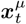, for *t* = 1 … *L*. Upon processing this sequence, the hidden activity ***h***^*µ*^ ∈ ℝ^*N*^, evolves according to

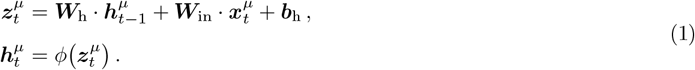

where ***W***_h_ ∈ ℝ^*N×N*^ is the recurrent weight matrix, ***W***_in_ ∈ ℝ^*N×α*^ is input weight matrix, ***b***_h_ ∈ ℝ^*N*^ are constant biases and *ϕ* is a non-linear function applied unit-wise. Here, we choose to work with rectified-linear units, i.e. *ϕ*(*z*) = max {0, *z*}. In the following, we will omit the sequence index unless necessary. All networks are trained via gradient-based optimization (Adam [37]) using backpropagation through time and performing mini-batch updates. Different tasks require different targets and loss functions, and the models used for different tasks only differ in the readout (see SI Sec. 4.2 for more details).

The central object of analysis throughout this work is the recurrent weight matrix ***W***_h_. We ask how its structure changes with learning, what that structure implies about the geometry of population activity, and what it reveals about the circuit-level basis of abstract temporal generalization. As we show below, the answer is a consistent one: learning drives ***W***_h_ toward a low-rank organization, and it is this low-rank structure — not the size or density of the network — that is the mechanistic substrate of abstract representation.

### 1.1 RNNs learn to generalize abstract sequence structure

We first examine a sequence classification task in which the network is trained to classify sequences based on their underlying temporal structure (see SI Sec. 4.1 for details on the generative process for constructing sequences). For example, sequences such as `abbb` and `addd` are generated from the class `ABBB`, while `abba` and `cddc` belong to the class `ABBA` (Fig. 1A). The network receives a label only at the end of the sequence, and the loss function is the cross-entropy between the output class probabilities generated by the network and the ground truth (see SI Sec. 4.2). In the remainder of the main manuscript, we focused on networks initialized in the “lazy” regime [38], where initial weights prior to training are drawn from a uniform distribution with a standard deviation of 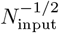, we have replicated all notable results in the “rich” regime in Fig. 10.

**Figure 1:**
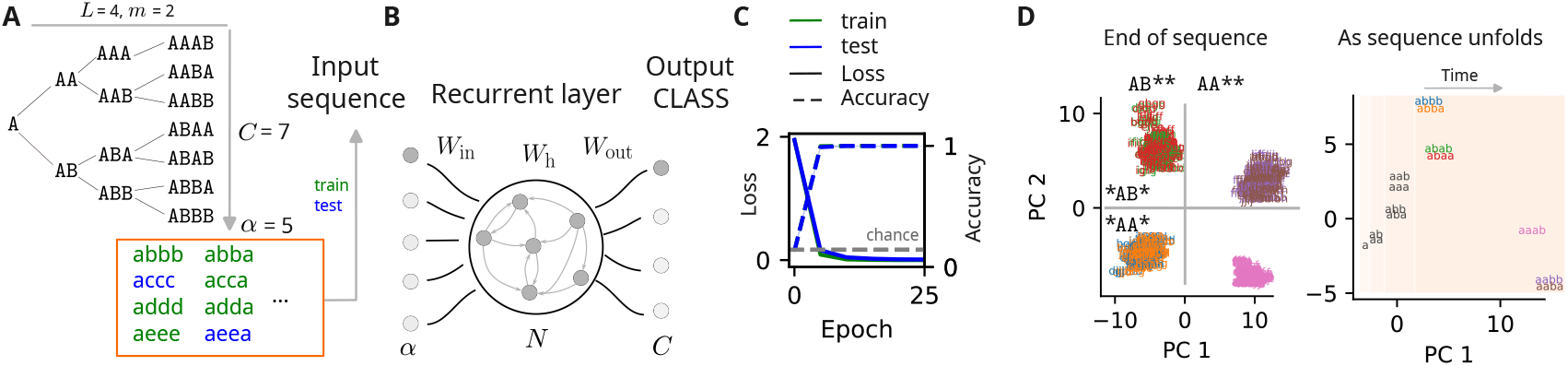
Emergence of abstract sequence representations in recurrent networks. **A**. Sequences are generated from a binary branching tree (example shown for sequence length *L* = 4, and *m* = 2 unique letters, corresponding to the number of branches from the root), yielding *C* = 7 abstract classes. Each class is instantiated by replacing the *m* unique letters with symbols drawn from an alphabet (of length *α* = 5). **B**. An RNN processes sequences one token at a time and is trained to classify them according to their abstract structure, receiving a class label only at the end. **C**. Training loss and accuracy across epochs (mean +/- s.d. across simulations). **D**. PCA of hidden activities at the end of training and end of sequence. Left: final hidden states cluster by abstract class and are linearly separable according to transition to same/different letter. Right: trajectories of hidden activity for one example sequence per class (indicated in different colors) plotted for each time-point as the sequence unfolds. PC2 and 3 capture branching dynamics that reflect the hierarchical structure of the generative tree as the sequence unfolds.

We find that sufficiently large networks achieve near zero loss on both training and test sets, indicating strong generalization performance (Fig. 1C and Fig. 6). Also, the number of classes that can be reliably generalized scales with network size, with larger networks required for generalizing a larger number of classes (Fig. 6, right).

In order to better understand the types of representations supporting generalization, we applied Principal Component Analysis (PCA) to the hidden activities at the end of the sequence. For sequences of length *L* = 4 and number of classes *C* = 4, the top three principal components account for ∼ 90% of the variance, revealing a highly structured, low-dimensional geometry. Moreover, the hidden activities strongly cluster by class, and are linearly separable according to transition types: whether the sequence transitions to the same or a different letter (Fig. 1D, left). When we projected the hidden activities onto the same reduced space, as the sequences unfold, we observed a branching pattern that mirrors the hierarchical tree structure used to generate the sequences (Fig. 1D, right). These results suggest that the RNN constructs internal representations that compress and organize sequence information in a way that reflects the abstract structure of the task. Next, we investigate the geometry of these unfolding representations, and the recurrent mechanism that supports it.

#### 1.1.1 Low-dimensional dynamics reflects low-rank recurrent connectivity

The compact, branching geometry of the population state described above implies that the recurrent weight matrix ***W***_h_ cannot be a dense, high-dimensional object — it must have structured, low-rank organization. To characterize this, we track how the dimensionality of population activity evolves over the course of each sequence, measuring it as the number of singular vectors of the population activity matrix required to capture at least 90% of the mean squared activity. Population activity is consistently low-dimensional relative to the input space, and dimensionality decreases progressively toward the end of the sequence (Fig. 2A). The trajectory is class-dependent: dimensionality rises transiently when a token arrives that differs from the previous one — momentarily expanding the representational space as a new branch of the tree is opened — but then collapses again as the sequence converges toward a single classifiable pattern.

**Figure 2:**
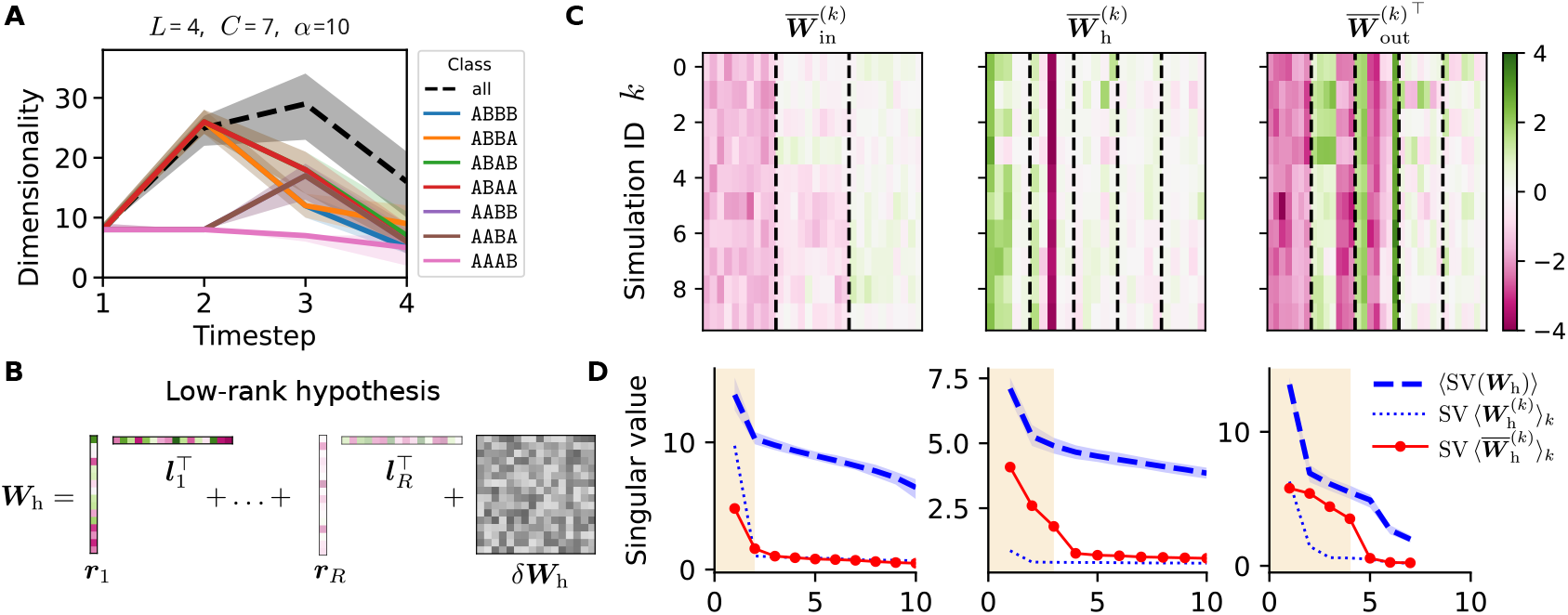
Low-dimensional dynamics and low-rank connectivity in trained recurrent networks. **A**. Dimensionality of hidden activity over time, measured as the number of singular vectors capturing 90% of the mean squared activity (medians +/- 5th-95th percentiles). Dimensionality is class-dependent; typically rising mid-sequence and decreasing toward the end. **B**. Recurrent connectivity matrix shown as a sum of a structured low-rank component (colored) and a random component (gray). **C**. Input, recurrent and output weight matrices from multiple simulations shown in a common rotated coordinate frame (see Eqs. 16). Each row corresponds to one simulation; matrices are flattened by concatenating rows (input/recurrent) or columns (output transpose). Only the dominant modes are shown (3 for input (left), 5 by 5 for recurrent (middle) and 5 for output (right) weights). Note the consistent structure across simulations. **D**. Low-rank structure extracted by averaging aligned weight matrices across simulations. Spectra of recurrent matrices are similar across simulations (dashed blue line: mean of SVs of the weight matrices from individual simulations +/- 5 − 95% percentiles bands). Spectra of the average recurrent weights before and after change of basis (dotted blue vs red) differ only in their structured components (shaded red region, used to infer the rank of the structured part). Left: spectrum of input weights. Middle: recurrent where *R* = 3. Right: output.

This progressive dimensionality reduction is the signature of low-rank recurrent connectivity [27]. Formally, we decompose the recurrent weight matrix as

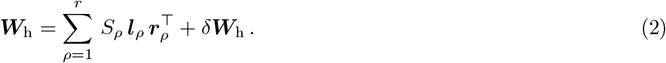

The first term is a structured, low-rank component — a sum of *r* outer products, each defined by a pair of left and right singular vectors ***l***_*ρ*_, ***r***_*ρ*_ with singular value *S*_*ρ*_ — and the second term *δ****W***_h_ is a residual reflecting variability from random initialization and the stochasticity of gradient descent (Fig. 2B). Biologically, each rank-one term 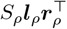 could represent a structured cell-type-specific projection: a population of neurons with input selectivity aligned with ***r***_*ρ*_ projecting to a population with output tuning aligned with ***l***_*ρ*_. The residual *δ****W***_h_ represents the background of unstructured, task-irrelevant synaptic variation that coexists with this structured signal.

To estimate the rank *R* of the structured component, we exploit the fact that networks trained on the same task with different random initializations should learn the same low-rank structure, differing only in a global rotation of the activity space. We therefore align the weight matrices of an ensemble of independently trained networks into a common coordinate frame (see Methods for the alignment procedure), and compare the singular value spectra before and after alignment. After alignment, averaging across networks preserves the common low-rank structure while canceling the random residual; before alignment, both are averaged away together. The rank *R* is identified as the number of singular values that survive this averaging — the “elbow” in the spectrum of the aligned average (Fig. 2D). For the binary tree tasks studied here, *R* = 3: just three structured outer products in the connectivity matrix account for the network’s capacity to represent and classify abstract sequence patterns. This is remarkably compact given that the network has *N* = 160 units and must distinguish up to *C* = 7 abstract classes.

Critically, the structured component is consistent across independently trained networks: when aligned into a common reference frame, the dominant modes of the input, recurrent, and output weight matrices are nearly identical across simulations (Fig. 2C). This consistency confirms that the low-rank structure is not an artifact of a particular initialization but reflects a universal solution to the task — the same abstract scaffold is discovered regardless of where optimization starts.

### 1.2 Low-rank recurrent connectivity gives rise to hierarchical relational geometry

Having established that classification drives the emergence of low-rank recurrent connectivity, we now ask: what is the functional relationship between this low-rank structure and the abstract relational representations observed in the population state? Specifically, do the dominant singular components of ***W***_h_ encode the same/different transition structure that defines the abstract classes?

To address this, we first examine the pattern of similarity among population states across sequences. If the network has learned to represent abstract class identity rather than token identity, sequences belonging to the same class should have similar population states — even when composed of entirely different tokens. We compute the cosine similarity between population state vectors across all pairs of sequences as the sequence unfolds (Fig. 3A). A clear structure emerges: from as early as the second token, the similarity matrix organizes into blocks that reflect transition type rather than token identity. Sequences sharing a “same” transition (`AA`-type) become more similar to each other than to sequences with a “different” transition (`AB`-type), independently of what specific tokens were used. At later timepoints, within each block, finer structure appears: sub-blocks form among sequences that also share their most recent transition. By the end of the sequence, the dominant organization reflects the *earliest* transitions — sequences that diverged at the first branching point of the tree are the most dissimilar, while those sharing the initial branch remain clustered together.

**Figure 3:**
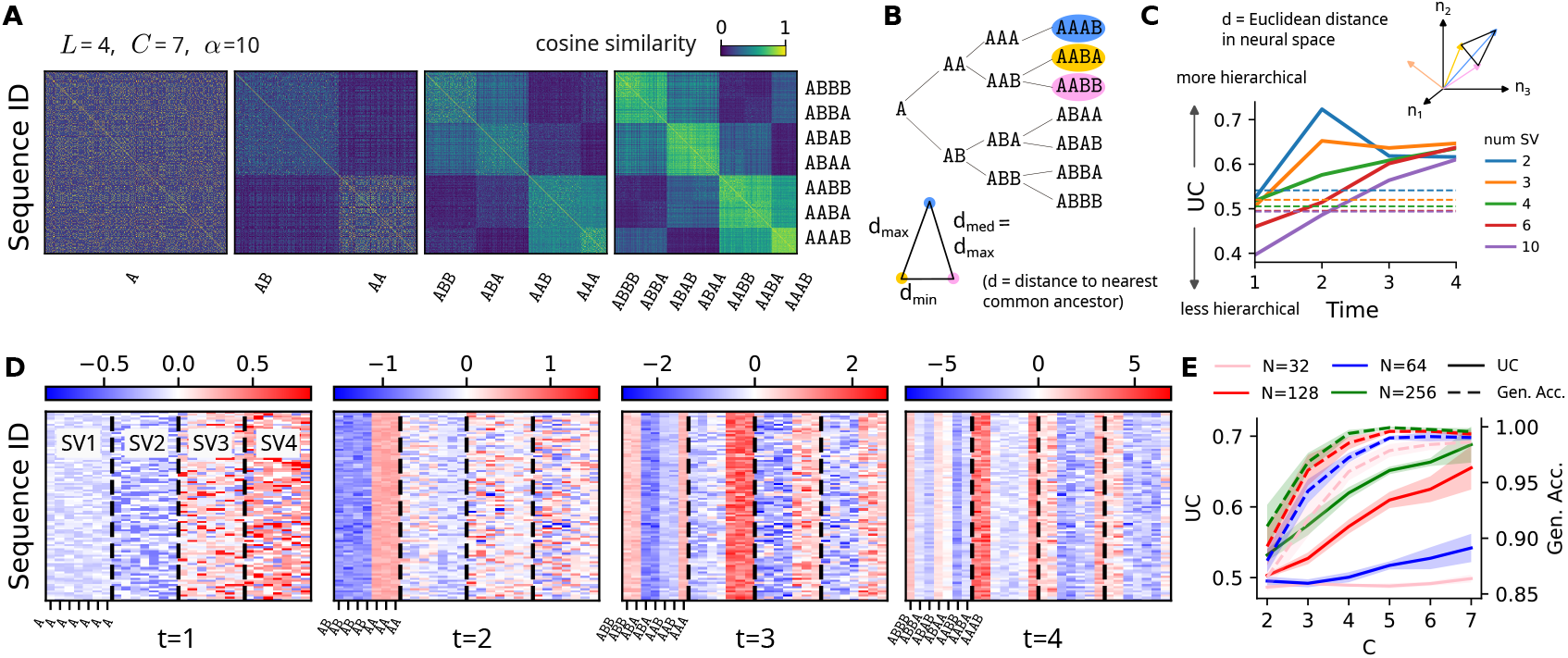
Hierarchical structure of hidden representations. **A**. Cosine similarity between hidden activities for all pairs of training sequences (test sequences show similar trends). As sequences unfold, the similarity matrix reveals a quasi-ultrametric organization reflecting the hierarchical structure of the sequences. **B**. Left: Hierarchical tree of abstract sequence classes. In an ultrametric space (bottom), distances between triplets of representations, as measured by their distance to their nearest common ancestor are restricted to equilateral or isosceles triangles with two long sides, disallowing isosceles with two short sides. A perfect hierarchical tree lies in an ultrametric space. **C**. Pairwise Euclidean distances between neural representations (top) yield ultrametric content (UC) values that approach–but deviate from–the ultrametric limit (bottom). UC over time for sequences of length *L* = 4. Solid lines: UC from projections onto singular vectors of the recurrent weights; four components capture the unfolding hierarchy. Dashed lines: UC within individual classes (baseline). **D**. Hidden activity projected onto the first four singular vectors of the recurrent connectivity. Rows: individual sequences; inner columns: class membership; outer columns (separated by dashed lines): the first four singular vectors. **E**. UC scores (left axis) and generalization accuracy (right axis) as a function of number of abstract classes. Higher UC scores correlates with higher generalization performance.

The resulting similarity structure approximates a binary branching tree in which sequences are organized chronologically according to their transition history (Fig. 1A). We quantify the degree to which population geometry conforms to this tree-like, hierarchical organization using the *ultrametric content* (UC) — a measure of how well a set of pairwise distances satisfies the ultrametric inequality [39, 40]. Intuitively, in a perfect ultrametric (tree-like) space, any three points form either an equilateral triangle or an isosceles triangle with the two long sides dominating; a high UC score indicates that the population geometry closely approximates the structure of a hierarchy (see Methods for formal definition). We find that UC increases monotonically as the sequence unfolds (Fig. 3C, solid lines), growing well beyond the within-class baseline (dashed lines) — indicating that the cross-class hierarchical structure, not just within-class clustering, becomes progressively more pronounced.

Crucially, this hierarchical structure is expressed in a compact, low-dimensional form that is directly linked to the recurrent connectivity: projecting the population activity onto just the top singular vectors of ***W***_h_ recovers most of the ultrametric structure observed in the full population state (Fig. 3C). Each successive singular component captures successive levels of the branching hierarchy (Fig. 3D): the first singular component separates sequences by their first transition, the second by their second, and so on. The low-rank connectivity is therefore not merely compressing activity into a smaller space — it is organizing that space according to the abstract relational structure of the task.

We note that the forward (chronological) tree is not the only hierarchical structure consistent with the population geometry. The cosine similarity matrix contains both strong diagonal blocks — reflecting early, forward transitions — and weaker off-diagonal blocks that reflect shared *recent* transitions, implying that the representations interpolate between a forward tree and a backward tree ordered from the last transition. This is not noise: it shows that the network retains information about the full transition history, and that early transitions are weighted more heavily in the final representation because the network processes sequences in temporal order.

Finally, we asked whether the quality of the abstract representation, as measured by UC, predicts the network’s capacity to generalize. For increasing numbers of abstract classes, we find a positive correlation between the mean UC score at the end of training and held-out generalization accuracy (Fig. 3E): networks that develop more strongly hierarchical population geometry generalize better. This demonstrates that the tree-like representational geometry is not merely a correlate of task performance — it is the mechanistic substrate of generalization. Having established this link, we next ask whether the low-rank component is causally necessary, and which specific component of the recurrent connectivity carries the critical computational role.

### 1.3 Perturbation experiments reveal functional role of dominant singular components

The analyses above establish that classification-trained networks spontaneously develop low-rank recurrent connectivity, and that this structure organizes population activity into a hierarchical, tree-like geometry. But are these structural components causally necessary for abstract generalization, or merely correlated with it? And which specific component does the critical computational work? We address these questions through two complementary perturbation approaches: constraining the rank of the connectivity during training, and selectively removing individual singular components from trained networks.

Because the low-rank structure and the random noise component of ***W***_h_ have comparable singular value magnitudes (Fig. 2D), the singular vectors of the full weight matrix do not cleanly isolate the structured components. We therefore first train networks in which the recurrent weights are explicitly constrained to rank *r* by parametrizing ***W***_h_ = ***AB***^⊤^, where ***A*** and ***B*** are ℝ^*N×r*^ matrices. This tests whether low-rank connectivity is *sufficient* for the task. Networks with *r* ≥ 3 ultimately match the generalization performance of unconstrained networks, though they learn more slowly (Fig. 4A). Networks with *r* = 1 or *r* = 2 show persistent, systematic failures: their confusion matrices reveal that they cannot distinguish abstract classes that differ in early transitions, collapsing distinct classes into indistinguishable groups (Fig. 4B–C).

**Figure 4:**
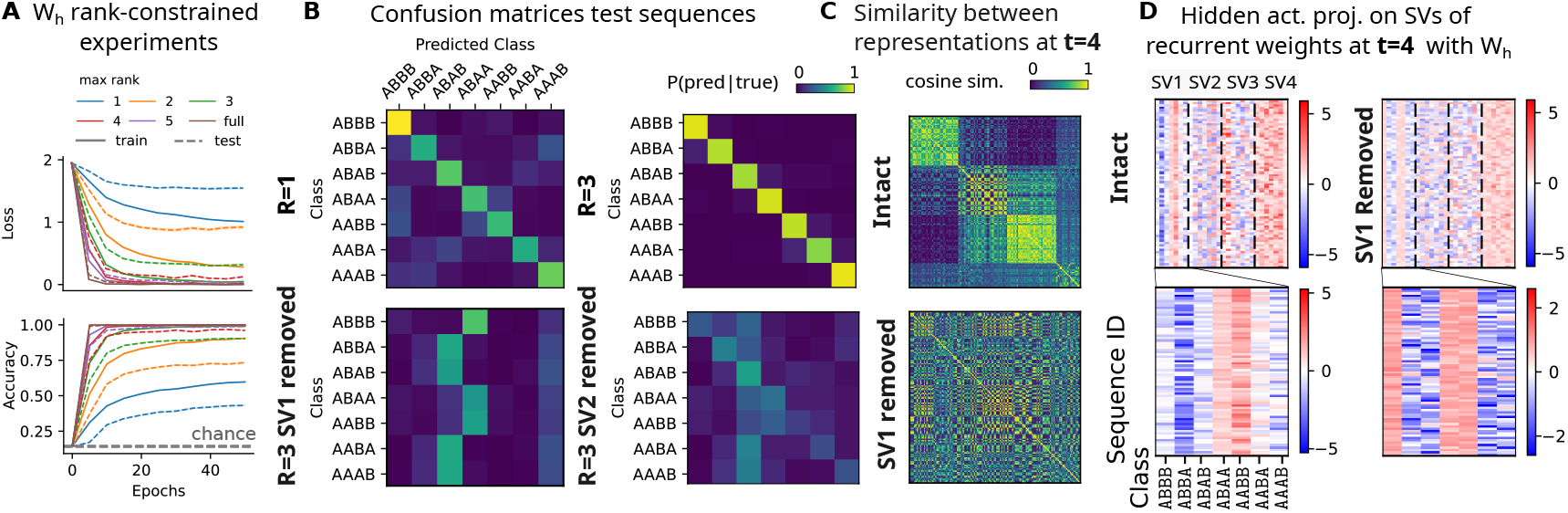
Perturbation of singular components reveals how low-rank dynamics support abstraction. **A**. We train networks with rank-constrained connectivity (ranks 1 − 5) and find that networks with low-rank recurrent connectivity can learn the task. **B**. Confusion matrices reveal the nature of errors. Perturbing a trained rank-3 recurrent matrix by removing individual singular components (bottom) show that eliminating the leading component (SV1) causes sequences to be classified almost solely by their last transition, “same” vs “different” (i.e. `AA` vs `AB`, collapsing many classes (bottom left). Removing SV2 yields milder distortions (bottom right). **C**. Cosine similarity matrices show that the structured clustering present in the intact network is disrupted when SV1 is removed. **D**. Projections of hidden activity onto the dominant (right) singular vectors illustrate the perturbation’s effect: with SV1 removed, the projection along that direction reflects only the most recent transition, while other projections carry little class information. In the intact network, SV1 (and to a lesser extent SV2-SV3) integrate transition information over time, supporting the formation of hierarchical, class-specific representations (see Fig. 9 for full time courses).

To interpret the functional role of the singular components, we performed *singular vector ablation* on trained networks with recurrent weights ***W***_h_. After training a rank-constrained network, we removed selected singular vectors from the recurrent weights, then simulated the network on all sequences, and computed its performance. The perturbed recurrent weight matrix where the *a*-th singular component has been removed would be 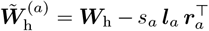, such that 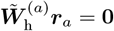 i.e. any activity along ***r***_*a*_ is not propagated to the next step.

Examining hidden activity projected onto the dominant singular vector reveals its functional meaning. At *t* = 2, projections along the first singular vector split cleanly into two blocks reflecting same/different transitions (`AA/BB` vs. `AB/BA`, Fig. 9). At subsequent timesteps, each block subdivides according to the *most recent* transition. By *t* = 4, this yields perfect decoding of abstract classes: the projection carries integrated information about the full transition history, hence the hierarchical generative tree (Fig. 4D left and Fig. 9B). However, when the dominant singular component is removed, this integrated structure collapses. The projection onto ***r***_1_ now reflects *only the very last transition*, and information about earlier transitions is lost (Fig. 4D, right and Fig. 9B). Therefore, the projected activity can only discern between sequences of type `**AB` vs `**AA`, but not *within* these groups. Correspondingly, the cosine similarity matrix loses its hierarchical block structure (Fig. 4C), and the confusion matrix shows that sequences are grouped solely by their *most recent* transition (Fig. 4B). Removing the second singular component produces only mild degradation, indicating that the dominant singular mode plays a special role in integrating sequential transition information over time.

More generally, ***r***_1_ retains a consistent functional meaning across the entire sequence: at every timestep, the projection of hidden activity onto ***r***_1_ selectively encodes the most recent same/different transition (Fig. 9). Removing the dominant singular component prevents this transition information from being propagated forward in time, so the network can no longer accumulate a history of transitions. As a result, the hierarchical, tree-like representation built from integrating successive transitions is disrupted, and only the most recent transition is retained.

Finally, in full-rank networks, ablating the dominant component produces qualitatively similar effects (Fig. 9), though somewhat attenuated due to residual noise components. The perturbed network still represents the most recent transition, but this information fails to propagate through time, preventing accumulation of relational structure.

Together, these findings demonstrate that the leading singular components of the recurrent weight matrix implement temporal integration of relational (“same/different”) transitions, enabling the network to construct tree-like representations of abstract sequence structure.

## 2 Task objective determines whether low-rank structure and abstract representations emerge

The results above raise a natural question: is low-rank recurrent organization a generic property of RNNs that process structured sequences, or is it specific to the classification objective? To answer this, we trained the same architecture on a next-token prediction task, in which the network must generate the next element at each timestep rather than wait for a global label at the end. Although classification and prediction share many computational demands — encoding time-varying inputs, maintaining memory across timesteps, and using knowledge of the sequence to constrain outputs — they differ in one critical respect: prediction can in principle be solved using only local, one-step transition statistics, without ever integrating information across the full sequence. Classification, by contrast, requires global integration: the label is only available at the end, so the network must carry a compressed summary of the entire transition history forward in time.

A practical challenge for training the prediction network was constructing sequences with unambiguous continuations. Because many partial sequences are consistent with multiple abstract classes, an unconstrained prediction network tends to memorize training-set continuations rather than generalizing. We therefore generated unambiguous prediction sequences by pruning the binary tree, retaining only a single valid continuation path for each starting cue (Fig. 5A). This ensures that each prediction target is uniquely determined by the cue, creating a fair comparison with the classification task.

**Figure 5:**
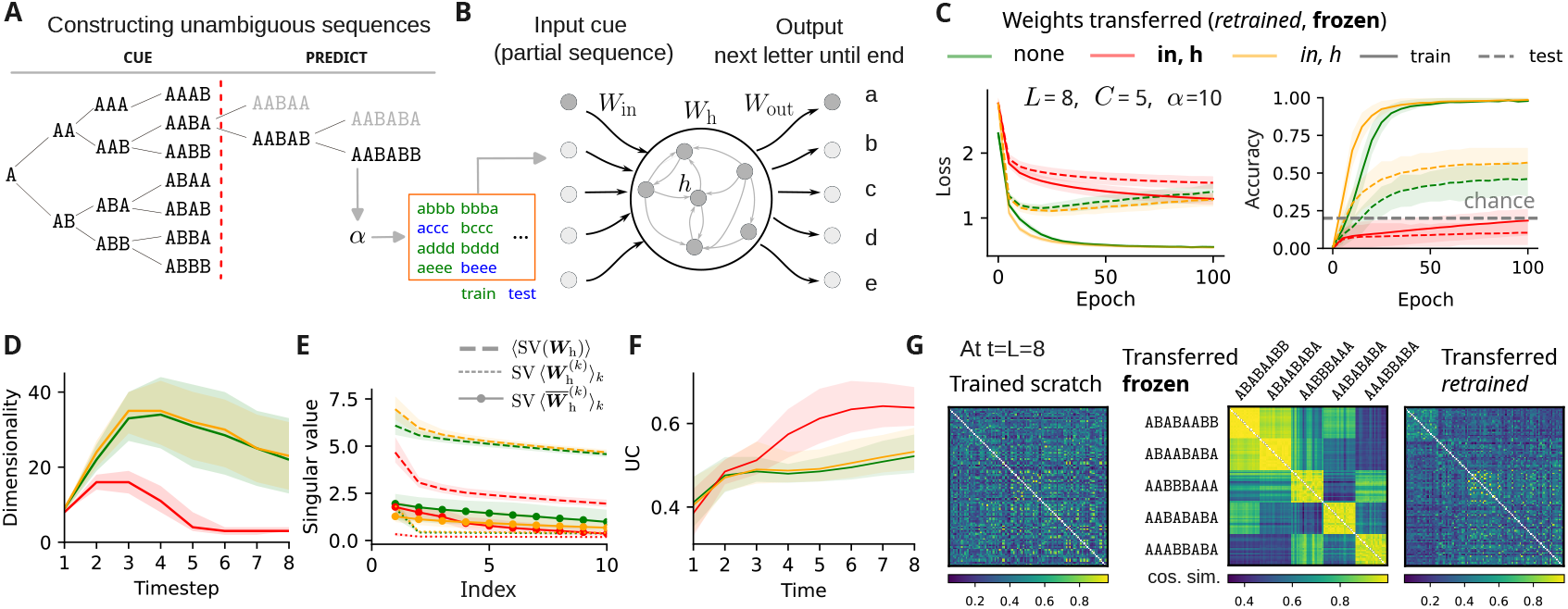
Transfer from classification improves prediction learning. **A**. To construct unambiguous sequences for the prediction task, we apply extreme pruning to a binary tree (light gray), retaining only one path per cue (black). The red dashed line marks the boundary between the cue (input) and the continuation to be predicted. Unique tokens are then replaced by letters drawn from an alphabet of size *α*. **B**. Left: a standard RNN is trained from scratch to predict each next token in the continuation sequence. Right: prediction networks are either initialized with weights transferred from a classification-trained RNN and then retrained (italic labels), or evaluated with transferred recurrent weights kept fixed (bold). **C**. Loss (left) and accuracy (right) for prediction networks under three initializations: random (green), transferred recurrent weights kept frozen (red; only output weights trained), and transferred recurrent weights used as initialization and then jointly trained (orange). Means and standard deviations shown across simulations. Only the latter condition accelerates memorization and improves generalization. To reduce overfitting, training was stopped at epoch 25 in all cases. **D**. Dimensionality of hidden activity over time, measured as the number of singular vectors capturing 90% of the mean squared activity (Median +/- 5 − 95% percentile bands). Dimensionality decreases mainly when transferred recurrent weights are frozen (red); networks trained from scratch (green) or fine-tuned from classification (orange) maintain higher dimensionality. **E**. Spectra of recurrent matrices (averaged over simulations, dashed), and spectra of their averages before and after change of basis (dotted vs solid). Only frozen transferred weights (red) yield a clear structured low-rank component (*R* = 3). Prediction networks trained from scratch (green) show no low-rank structure; those initialized from classification (orange) also lack clear low-rank structure, though their spectrum is less flat than randomly initialized networks. **F**. Ultrametric content (UC) of hidden representations over time for each initialization condition (as in Fig. 3C, mean UC +/- 5 − 95% percentile bands). **G**. Cosine similarity matrices between hidden representations for each initialization condition. Left: prediction net trained from scratch. Middle: weights transferred from classification to prediction and frozen. Right: weights transferred from classification and then retrained. Only the transfer-and-freeze condition preserves the hierarchical, low-dimensional representations.

Despite seeing the same sequences, prediction networks behave fundamentally differently from classification networks. They achieve strong memorization of training sequences but generalize poorly to held-out ones (Fig. 7). More tellingly, their internal dynamics differ at the circuit level: population dimensionality remains high throughout the sequence rather than progressively compressing (Fig. 5D, green); the recurrent weights show no low-rank structure (Fig. 5E); and their representations lack hierarchical organization, as measured by UC (Fig. 5F). Low-rank dynamics do not therefore arise from processing structured sequences per se — they are specific to tasks that require global temporal integration, consistent with the idea that the task objective is the key variable shaping circuit organization [41, 42].

Having established that classification scaffolds low-rank structure while prediction does not, we asked whether the classification-trained scaffold could be transferred to bootstrap generalization in the prediction task. We initialized prediction networks with recurrent and input weights from a classification-trained model and tested two regimes: keeping the transferred weights frozen (only the readout is retrained), or using the transferred weights as an initialization for continued training. Freezing the weights provides no benefit — the frozen scaffold cannot be read out by a randomly initialized readout (Fig. 5C, red). Using the scaffold as an initialization, however, substantially accelerates learning of the test set and improves asymptotic generalization performance (Fig. 5C, orange). The benefit is primarily carried by the recurrent weights rather than the input weights (Fig. 8A), consistent with the idea that the abstract schema resides in the internal circuit organization — the recurrent connectivity — rather than in the input representations.

Critically, this improvement is specific to the abstract scaffold and not a generic effect of pretraining. An autoencoder trained to reconstruct the same sequences transfers weights that speed up memorization — a warm-start effect — but do not improve generalization (Fig. 8E–F). The dissociation between classification pretraining (which improves generalization) and autoencoder pretraining (which does not) demonstrates that the transfer benefit depends on the relational structure learned by the scaffold, not on statistical familiarity with the sequence distribution. Prior exposure to the same sequences is not enough; what matters is having learned to compress them into a hierarchical, low-rank internal representation.

## 3 Discussion

We showed that abstract temporal structure can emerge spontaneously in recurrent neural networks trained only on sequence-level labels. Without architectural biases or intermediate supervision, standard RNNs develop compact, low-dimensional internal representations that reflect the hierarchical structure of the input sequences and generalize across token identities. This abstraction is mechanistically supported by low-rank recurrent connectivity, which constrains population dynamics to a shared low-dimensional manifold aligned with the generative tree, and by a dominant recurrent mode whose causal role is to integrate relational transition information across time. Importantly, this geometry is not merely a byproduct of learning: networks that develop more strongly hierarchical, ultrametric representations generalize better. Together, these findings provide a circuit-level account of how recurrent architectures can internalize the latent relational structure of sequential experience and represent it in a form that supports flexible, identity-independent generalization.

### Low-rank connectivity as a synaptic substrate for abstract representation

A central contribution of this work is the identification of a *specific circuit property*—low-rank recurrent connectivity—that underlies the emergence of abstract relational representations. In the low-rank framework [27], the synaptic weight matrix is not a dense, high-dimensional object, but a sum of a small number of structured outer products, each implementing a distinct computational channel, superimposed on a random background. Our results show that learning to classify sequences by abstract pattern spontaneously drives recurrent connectivity toward this low-rank organization: for the binary-tree tasks studied here, the structured component has rank *R* ≈ 3, while the remainder of the weight matrix contributes primarily unstructured variability.

This observation admits a natural biological interpretation. In cortical and hippocampal circuits, the analog of a rank-one component may be a specific cell-type-to-cell-type projection: a population of neurons with shared input selectivity projecting coherently to a population with shared output tuning [27]. The effective rank of the circuit—the number of such structured components—determines the number of independent computations it can support. Our finding that rank 3 is sufficient to abstract a binary tree with 7 classes suggests that surprisingly few structured synaptic motifs may be needed to implement hierarchical relational coding. From a circuit-neuroscience perspective, this predicts that the core circuit supporting abstract sequence learning may be describable by a small number of cell-type-specific connectivity rules, and that disrupting one of these rules should have qualitatively different effects from disrupting an equivalent number of random synapses.

The dominant singular component deserves particular emphasis. Our ablation experiments show that it serves a specific and non-redundant role: at each timestep, it encodes the most recent same/different transition and propagates this information forward into the recurrent state, where it is accumulated across the sequence. Removing this component does not abolish sequence processing altogether—the network retains sensitivity to the most recent transition—but it selectively erases memory for *earlier* transitions, collapsing the hierarchical representational geometry into a flat, one-step code. This dissociation identifies a concrete circuit mechanism for integrating relational history over time. In biological circuits, the neurons most strongly aligned with this dominant mode—those whose activity is most strongly modulated by the component’s input and output patterns—would be especially important targets for perturbing abstract sequence representations.

### Task objective determines circuit organization

A striking result is that the same input statistics, processed by the same architecture, give rise to fundamentally different circuit organization depending on the task objective. Networks trained on next-token prediction—which requires only local, one-step forecasting—fail to develop low-rank recurrent structure or hierarchical population geometry, even though they process identical sequences. Lowrank structure and abstract representation emerge specifically when the task requires *global temporal integration*: maintaining a compressed summary of the full transition history and making a judgment that depends on the sequence as a whole rather than only on the most recent element.

This has direct implications for understanding what different brain regions may compute during sequence learning. If a circuit’s effective task objective is primarily local and predictive—as may often be the case for sensory cortex or cerebellar-like circuits—one would not expect abstract, low-rank organization to emerge. By contrast, circuits that integrate information across longer temporal windows, or that receive feedback at the resolution of whole events or episodes rather than individual stimuli, are natural candidates for the kind of low-rank, abstract structure described here. This interpretation is consistent with the established roles of hippocampus and prefrontal cortex in temporal integration and rule-based generalization [5, 7]: these regions are anatomically and functionally positioned to support the longer-timescale computations required for schema formation.

### Schema reusability and the neural basis of transfer

Our transfer experiments reveal a further property of the low-rank scaffold: it is *reusable*. Initializing a prediction network with recurrent weights from a classification-trained network accelerates learning and improves generalization, and this effect is specific to the abstract scaffold rather than reproduced by autoencoder pretraining on the same data. The benefit arises primarily from transferring the recurrent weights rather than the input weights, consistent with the idea that the internal circuit organization—rather than the input representations—carries the abstract schema. This aligns naturally with schema-accelerated learning in the biological literature [9], where prior structured knowledge dramatically speeds the acquisition of new memories that conform to the same relational structure.

In the rodent schema experiments of Tse and colleagues, animals that had learned a set of flavor-place associations in a particular spatial context could acquire new associations in that same context within a single trial, whereas naive animals required weeks of training [9]. Our model suggests a mechanistic interpretation of such effects: prior learning builds a low-rank recurrent scaffold encoding the abstract relational structure of the environment, and new experiences that share that structure can be rapidly assimilated because the circuit already contains the motifs needed to represent them. The specificity of transfer in our model—classification-trained scaffolds help, generic pretraining does not—parallels the specificity of schema effects in memory, where prior knowledge accelerates learning of schema-consistent content but not content that is merely statistically similar without being relationally congruent [9, 43, 44].

The fact that *freezing* the transferred weights provides no benefit is also informative. It indicates that the scaffold does not directly encode the new task, but instead provides a favorable initialization from which learning can rapidly discover a suitable solution. This is consistent with viewing schemas as reusable *structural priors* on circuit dynamics rather than stored solutions, and may map onto the distinction between hippocampal contributions to rapid encoding and prefrontal contributions to long-term schema storage and deployment [45].

### Predictions for neural population activity

The low-rank framework generates concrete predictions for the geometry of population activity in circuits engaged in abstract sequence learning, which we are currently investigating in electrophysiological recordings from rodents.

First, population activity during sequence presentation should be *low-dimensional*, with dimensionality substantially lower than the number of recorded neurons and closer to the number of abstract classes than to the number of distinct tokens. Crucially, this low dimensionality should be structured: the leading principal components should carry class-diagnostic information and organize according to the hierarchical structure of the sequences, yielding branching, tree-like trajectories rather than arbitrary smooth manifolds.

Second, there should be a *dominant population axis* that selectively encodes the most recent same/different transition, independently of the identity of the tokens involved. This axis should also carry information about *earlier* transitions, reflecting its role as an integrator of relational history. Perturbing neurons most strongly aligned with this axis, while leaving the rest of the population intact, should selectively impair abstract class discrimination while sparing sensitivity to token identity.

Third, the degree to which population geometry is hierarchical and ultrametric—quantified here by ultrametric content—should *predict generalization performance* across animals, sessions, and circuit areas. Regions or sessions with higher ultrametric content should show better transfer to novel token instantiations of familiar abstract classes, providing a population-level neural correlate of abstract generalization rather than mere memorization.

Fourth, the distinction between classification and prediction tasks implies a corresponding distinction in neural circuit organization: circuits engaged in global sequence classification should exhibit lower-dimensional, more hierarchically organized population geometry than circuits performing only local prediction on the same stimuli. Thus, hippocampal population activity during rule-learning should be more ultrametric than during a matched next-step prediction task, even when the sensory input is identical.

### Relation to hippocampal and prefrontal function

The hippocampus is a natural candidate substrate for the low-rank scaffold described here. It is known to support the rapid formation of context-dependent and relational memories [5]; its population activity organizes into structured, low-dimensional representations that encode the relational geometry of experience [26]; and it appears to be required for the kinds of unsupervised, schema-dependent learning addressed by our model. Our recent recordings from hippocampal CA1 during passive exposure to structured auditory sequences show that dCA1 ensembles dynamically reorganize into subspaces that separately encode sensory features and abstract relational rules, and that this reorganization is causally dependent on hippocampal activity [4]. This is broadly consistent with the present model: a circuit with distinct low-rank components for identity encoding and relational structure would be expected to separate sensory and abstract dimensions of the task.

Prefrontal cortex likely plays a complementary role. Whereas hippocampus may construct the scaffold through experience-dependent plasticity, prefrontal circuits are thought to maintain and deploy abstract rules in a flexible, context-dependent manner [7, 25]. In our model, the critical substrate for transfer lies in the *recurrent* component of connectivity—the internal circuit organization—rather than in the input or output weights. This resonates with the idea that prefrontal-hippocampal interactions during consolidation progressively stabilize abstract structure in cortex, where it can later be flexibly deployed across contexts [45, 44].

### Limitations and future directions

Several limitations of the present work should be noted. First, we focus on algebraic patterns generated by deterministic binary trees and fixed sequence lengths; whether similar low-rank mechanisms extend to probabilistic grammars, variable-length sequences, or continuous-valued inputs remains open. The binary-tree structure provides a particularly clean instantiation of hierarchical relational organization, but real-world sequences encountered by animals—acoustic, olfactory, or social—are probabilistic, variable-length, and often hierarchically nested in more complex ways. Extending the framework to stochastic transition rules, in which the same/different signal is noisy, would require circuits to perform probabilistic inference over transition history rather than deterministic accumulation.

A related limitation is that the present model uses discrete, categorically distinct tokens. In many biologically relevant sequence-learning problems, stimuli vary along continuous dimensions and the relevant relational rule is defined over metric relationships among continuous values (for example, whether pitch is rising or falling). Extending the low-rank framework to such settings—where the dominant component would need to encode the sign of a continuous derivative rather than a categorical same/different relation—is an important direction for future work.

Perhaps the most important open question is how low-rank structure and abstract representations *emerge over learning*, rather than simply being present at convergence [46]. Our analyses focus on end-of-training representations; they do not characterize the trajectory from random initialization to structured low-rank connectivity. Understanding when the dominant singular component first becomes identifiable, whether its emergence is gradual or abrupt, and how it depends on curriculum and experience statistics would provide a richer account of the learning dynamics underlying schema formation.

Finally, the task objective plays a decisive role in our model: low-rank abstraction emerges only when the circuit is forced to integrate globally rather than operate locally. What determines whether a biological circuit is effectively performing global integration versus local prediction is unknown, and likely depends on neuromodulatory state, feedback connectivity, and the temporal statistics of behavioral experience. Understanding the conditions that favor one regime over the other—and thereby promote or prevent schema formation—remains an important question for future experimental and theoretical work.

## 4 Methods

### 4.1 Generating structured sequences

We construct sequences designed to exhibit specific regularities. Two key parameters define the sequence space: the sequence length *L*, and the number of distinct letters *m* used in each sequence, with *m < L*, to ensure repetition and thus induce “regularity” in the sequence. For example, consider *L* = 2 and *m* = 2: the only possible sequence is *AB*. With *L* = 3 and *m* = 2, possible patterns include *AAB, ABB* and *ABA*, each exhibiting a different structure of repetition and alternation–e.g. “same different” vs “different same”. These patterns define what we refer to as *algebraic patterns* [14], or abstract temporal schemas. The number of such abstract patterns correspond to the number of ways to partition a sequence of *L* items into groups of at least one element is given by

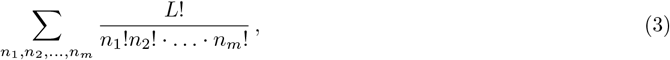

where the sum over *n*_1_, …, *n*_*m*_ is over partitions of *L*, i.e. they satisfy the constraint 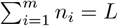.

To instantiate each abstract pattern with concrete sequences, or tokens, we select *m* distinct letters from a given alphabet of size *α*. The number of such choices is

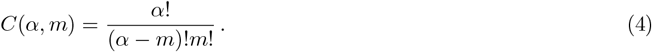

Thus, we produce *p* instances of the same sequence type, given by the multiplication of the two expressions above

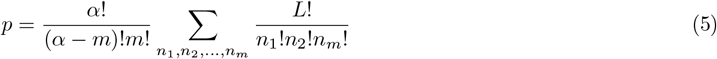

These sequences constitute a fraction of the total number of possible sequence configurations of length *L* given by *α*^*L*^. We use these structured sequences to train our model. In this manuscript, we used 80% of the sequences (chosen randomly) to train the models, and the remaining 20% to test the models. Choosing sequences randomly meant that networks were highly likely to have seen all the letters used in the data, but never in the configurations present in the test data.

#### Sequences produced using a binary branching tree

In the special case of *m* = 2, this generative procedure defines a binary branching tree: each additional token in the sequence corresponds to a binary split (of an existing node), producing a hierarchical structure over the abstract classes (Fig. 1A, main text). Throughout the main text, we focus on this case to study how networks learn and represent such tree-structured temporal regularities.

#### Generating unambiguous sequences for testing generalization in prediction networks

As mentioned in the main text, sequences generated from terminal nodes of the full binary branching tree naturally contain overlapping elements across classes. This overlap poses a challenge for evaluating generalization in the prediction task, which requires prompting the network with a partial sequence (of length *L*_*cue*_ *< L*), and having it recursively predict the remaining *L*_*retr*_ = *L* − *L*_*cue*_. This meant that a partial cue could theoretically lead to multiple different retrieval options, leading to ambiguous sequences. When ambiguous cues were used, the prediction network often defaulted to generating familiar training-set continuations, and was unlikely to retrieve sequences in the test set. This made it difficult to assess its ability to generalize to unseen sequences. To address this, we designed a procedure to construct unambiguous classes for testing prediction performance. Specifically, we generated cues of length *L*_*cue*_ using the full tree structure, and then appended unique combinations of *m* letters beyond that point, effectively pruning all but one continuation path from the node *L*_*cue*_ onwards. Conceptually, this amounts to constructing a tree of depth *L*, from which all-but-one branches from node *L*_*cue*_ to the terminal nodes *L* of the tree are pruned (Fig. 4A, main text). This procedure meant that for a specific choice of *L*_*cue*_ and *L*, many different combinations of classes are possible, from which we sample *C* at a time (Fig. 4A of main text, black sequences sampled, gray sequences not sampled). To make sure that any effect that we observe is not due to a specific choice of this “class combination (*CC*)”, we run many different simulations with different samplings, with a maximum of *CC* = 20 unique class combinations, when available.

### 4.2 Neural Network Models

#### Classification model

In this task, the output of the RNN model is the probability distribution ***p*** over *C* classes, therefore in the (*C* − 1)-dimensional simplex Δ^*C*−1^. We choose to minimize a cross-entropy (CE) function

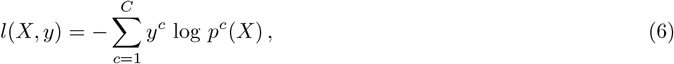

where *y*^*c*^ is the target probability that the sequence *X* belongs to class *c* (1 if *c* equals the true class, 0 otherwise). The output is computed after the *L*-th (last) element in the sequence *X*:

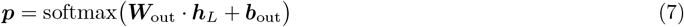

where ***W***_out_ ∈ ℝ^*C×N*^ is the output weight matrix, i.e.

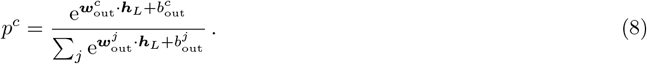

where 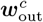 and 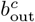 are the *c*-th row of ***W***_out_ and the *c*-th component of ***b***_out_, respectively.

#### Prediction model

In the prediction task, the output is a probability distribution over the letter in the alphabet, ***p*** ∈ Δ^*α*−1^. Here the objective is to predict the next letter in the sequence. Given a starting cue letter, the network outputs the next letter, and this next letter then serves as a cue for the prediction of the next letter. Therefore the loss function that we minimize is the cross entropy between sequence elements at time *t* and those at time *t* + 1.

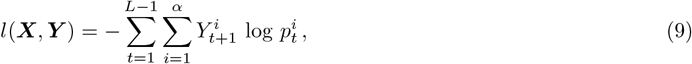

where 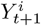 is the next letter in the sequence, and 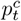 is probability distribution of the next letter as predicted by the network, given by

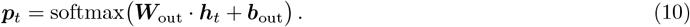

where ***W***_out_ ∈ ℝ^*α×N*^.

#### Reconstruction model

For this task, we consider an autoencoder architecture composed of an RNN encoder and an RNN decoder, with the same number of recurrent units, *N*.

At the end of the input sequence, a fully connected linear layer maps the recurrent activity of the encoder to an *n*-dimensional latent space.

The latent activity is then fed as a constant input to the decoder RNN through a linear. All the weights in the network are updated via batch gradient descent through BPTT.

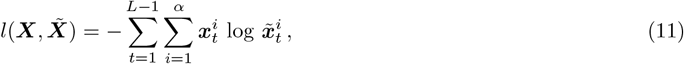

where 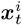 is the input sequence (and the target), 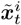 is the reconstructed sequence, given by

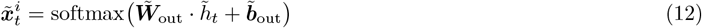

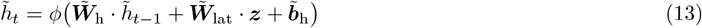

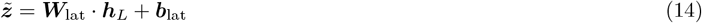

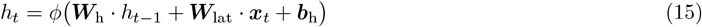

with ***h***_0_ = **0** and 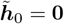 and where 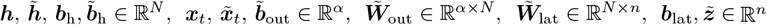, and ***W***_lat_ ∈ ℝ^*n×N*^.

### 4.3 Estimating the rank of the structured connectivity component

Because networks trained from different random initializations learn the same low-rank structure but expressed in different coordinate frames, the structured component cannot be recovered simply by averaging weight matrices across runs — the random relative orientations cancel it. We instead align the weight matrices of an ensemble of *K* independently trained networks into a common coordinate frame before averaging.

The alignment is based on the population activity at the end of the sequence. For each network *k*, we collect the *p* × *N* matrix ***H***^(*k*)^ whose rows are the final hidden state vectors 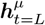 for all training sequences *µ*. We apply SVD to each activity matrix, 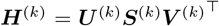, and use the right singular vector matrix ***V*** ^(*k*)^ to define a network-specific rotation 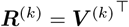. We then transform each network’s weight matrices into this common frame:

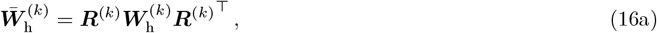

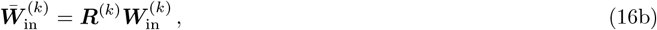

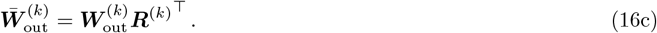

In the aligned frame, the structured low-rank component is consistently oriented across networks, so it survives averaging: 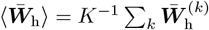 retains the common structured part. By contrast, in the original unaligned frame, the structured component is averaged away along with the noise: 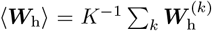 retains only the mean, which is near zero. The rank *R* of the structured component is estimated as the number of singular values of 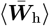 that exceed those of ⟨***W***_h_⟩ — identified as the “elbow” in the spectrum of the aligned average (Fig. 2D). The alignment procedure is exact for linear networks and holds approximately for ReLU networks, as verified by the consistency of aligned weight matrices across simulations (Fig. 2C).

### 4.4 Ultrametric content

Given a set of *p* representations 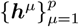, we first compute the distance between any of the pairs ***h***^*µ*^, ***h***^*ν*^. Fig. 3B of main text shows a graphical depiction of sequences constructed using the binary branching process. Within this tree, the distance between any two terminal nodes (shown in color) can be defined as the distance to the nearest common ancestor. In order to compute the distance between pairs of real-valued neural representations, we simply compute the Euclidean distance between vectors of neural activity at any time-point (Fig. 3C, main text, top). We then take triplets of representations, (***h***^*µ*^, ***h***^*ν*^, ***h***^*ρ*^). Denoting by 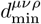 the edge of minimal length in the corresponding triangle, 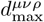 the edge of maximal length and 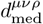 the edge of intermediate length, the ultrametric content [39] is defined as

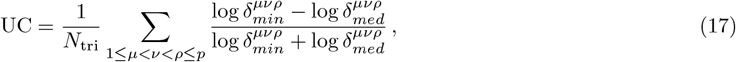

where *δ*_*min*_ = *d*_min_*/d*_max_ and *δ*_*med*_ = *d*_med_*/d*_max_.

In Fig. 3C of main text, we compute the UC for the set of representations corresponding to all sequences (hidden activity at any given time-point, solid lines). As a control, to verify that the tree structure emerges due to the configuration of classes relative to one another, we also compute the UC at the end of the sequences, restricted to triplets belonging to the same class. It can be seen that towards the end of the sequence, as the class membership becomes apparent, the UC computed over all representations is above the UC values computed for triplets restricted to individual classes (0.5, Fig. 3C, main text).

If we plot *δ*_*min*_ = *f*(*δ*_*med*_), triplets that satisfy the triangular inequality lie above the line *δ*_*min*_ = 1 − *δ*_*med*_, while triplets that satisfy the ultra-metric inequality lie on the vertical line where *δ*_*med*_ = 1. Triplets that are equilateral triangles lie at the point *δ*_*min*_ = *δ*_*med*_ = 1.

Eq. 17 gives a measure of the overall closeness of the cloud of triplets to the fully ultra-metric limit. This quantity ranges from 0 (all triplets forming isosceles triangles with two short sides) to 1 (for a fully ultra-metric set: equilateral triangles and isosceles triangles with two long sides).

## Acknowledgements

We thank members of the Akrami lab and other SWC laboratories for insightful discussions and advice. This work was supported by core funding to the Sainsbury Wellcome Centre from Wellcome (219627/Z/19/Z) and the Gatsby Charitable Foundation (GAT3755), and by UK Research and Innovation (EP/Z000599/1) to A.A.

## 5 Supplementary Material

### 5.1 Phase diagrams

To generate the phase diagrams for both the classification network (Fig. 6) and the prediction network (Fig. 7) where we vary the number of classes *C* and the length of the sequences *L*, we used the pruning procedure to construct unambiguous sequences. This also allowed us to run transfer experiments (Fig. 8).

**Figure 6:**
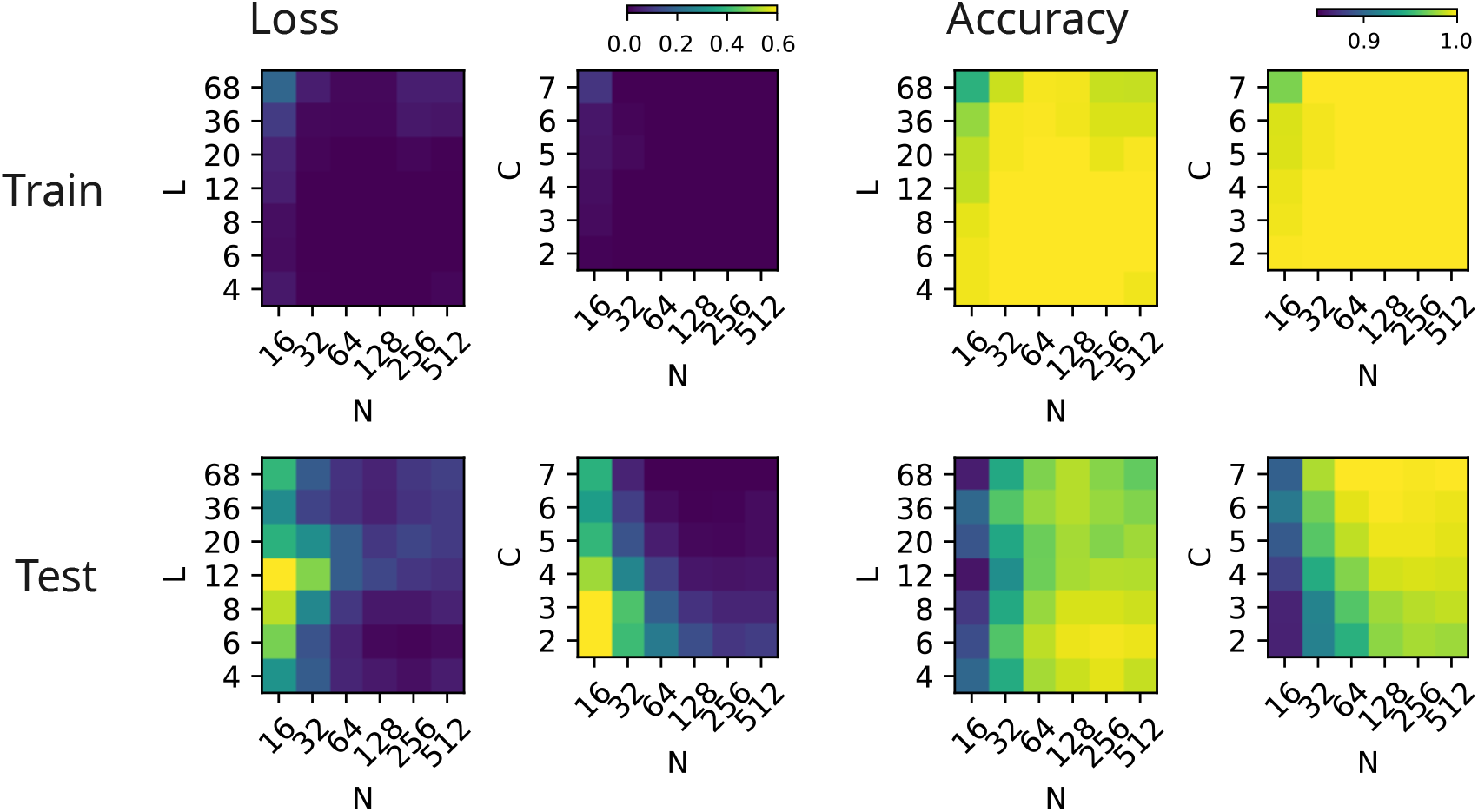
Classification Task. Phase diagrams showing train/test loss and accuracy as functions of network size *N* (x-axis) and either sequence length *L* or number of classes *C* (y-axes). When *L* is varied, the number of classes is fixed to *C* = 4. When *C* is varied, the sequence length is fixed to *L* = 7.

**Figure 7:**
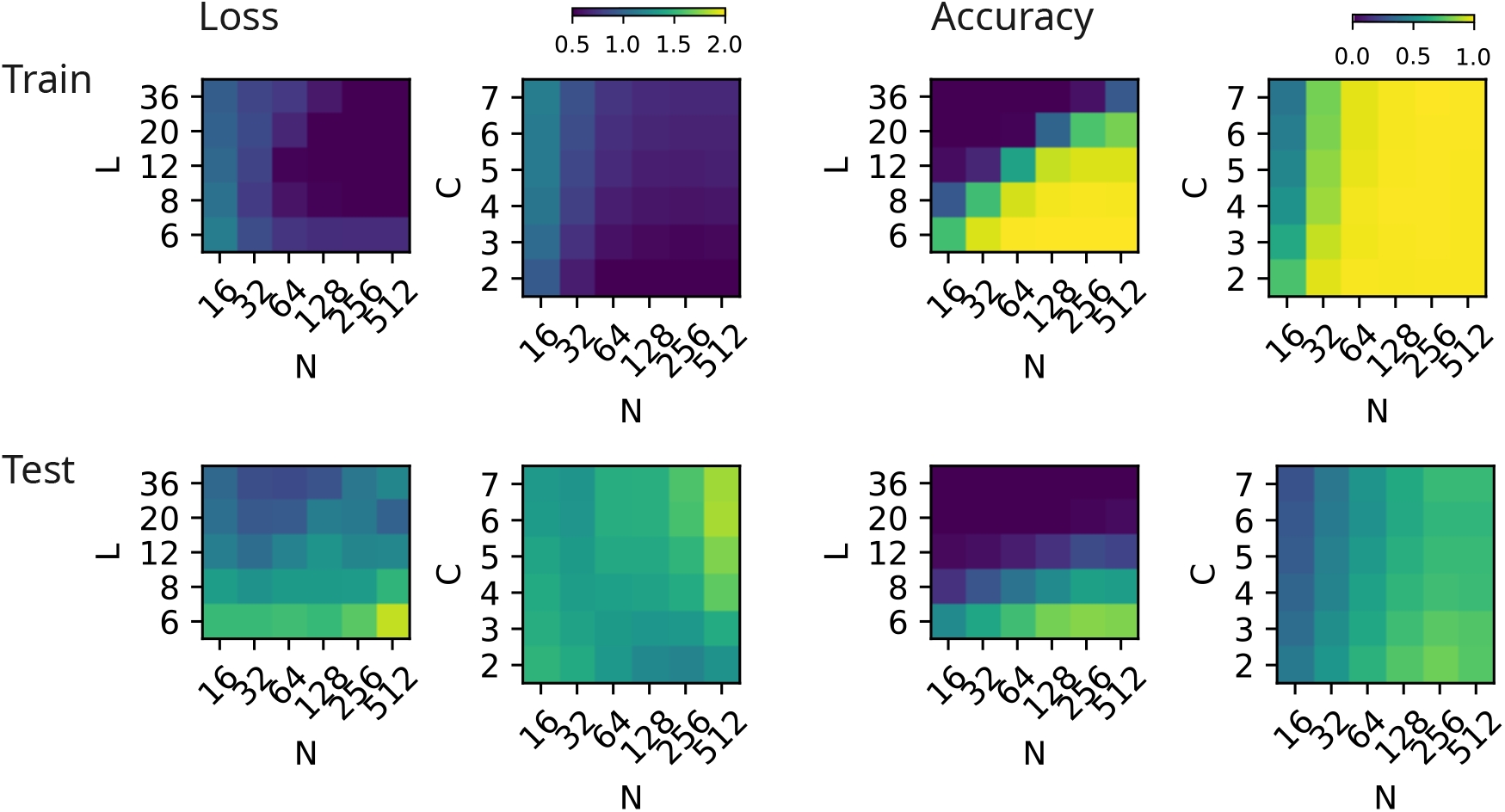
Prediction Task. Phase diagrams showing train/test loss and accuracy as functions of network size *N* (x-axis) and either sequence length *L* or number of classes *C* (y-axes). Networks reliably memorize training sequences, but show poor generalization to held-out sequences. Longer sequences and more classes require larger networks.

**Figure 8:**
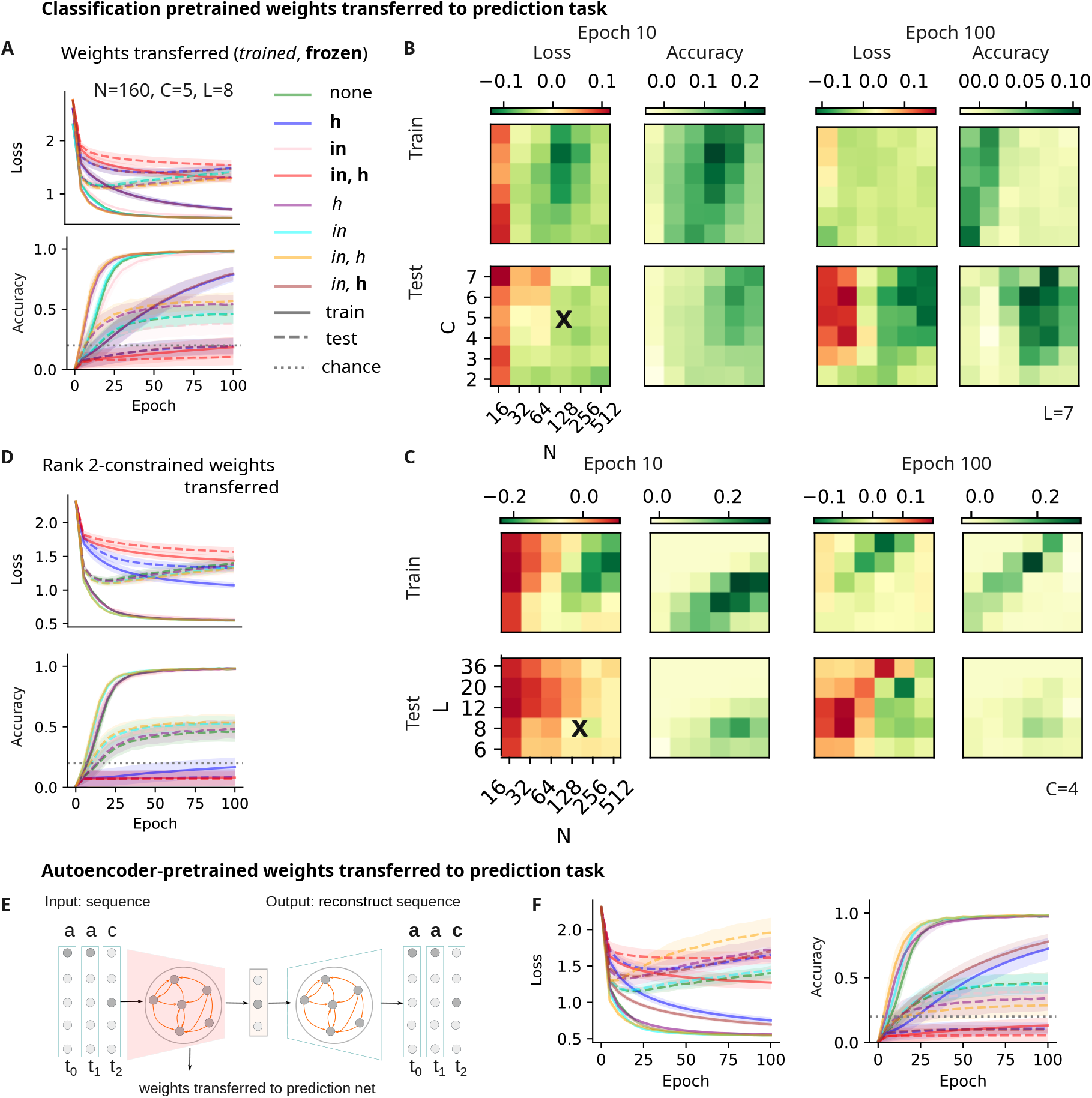
The effect of classification and reconstruction pretraining on prediction tasks. **A**. Loss (top) and accuracy (bottom) of prediction networks under three different initialization protocols: random initialization in the lazy regime (green), parameters from the classification pretraining transferred and kept frozen (blue, pink and red), and continued training (purple, cyan and orange). Finally, brown indicates frozen recurrent weights and retrained input weights. Lines indicate the mean across simulations; shaded areas show standard deviations. **B**. Difference in loss and accuracy between transfer-initialized and randomly initialized prediction networks, as a function of network size *N* (x-axis) and number of classes *C* (y-axis), for *L* = 7. These phase diagrams are plotted for early training (epoch 10) and late training (epoch 100). Green areas indicate improved performance through transfer. The black cross marks parameter configuration shown in panel **A. C**. Same as **B** but with sequence length *L* on the y-axis, for fixed number of classes *C* = 4. **D**. We pretrained rank-constrained networks on the classification task and then transfer the weights onto the prediction network following the same protocol as in **A. E**. We perform the same transfer experiments by pre-training on a reconstruction task, with the RNN as the encoder of an auto-encoder architecture. **F**. The loss and activity learning curves show that transfer makes it faster to learn the training data, but it does not lead to an improvement in the generalization performance.

**Figure 9:**
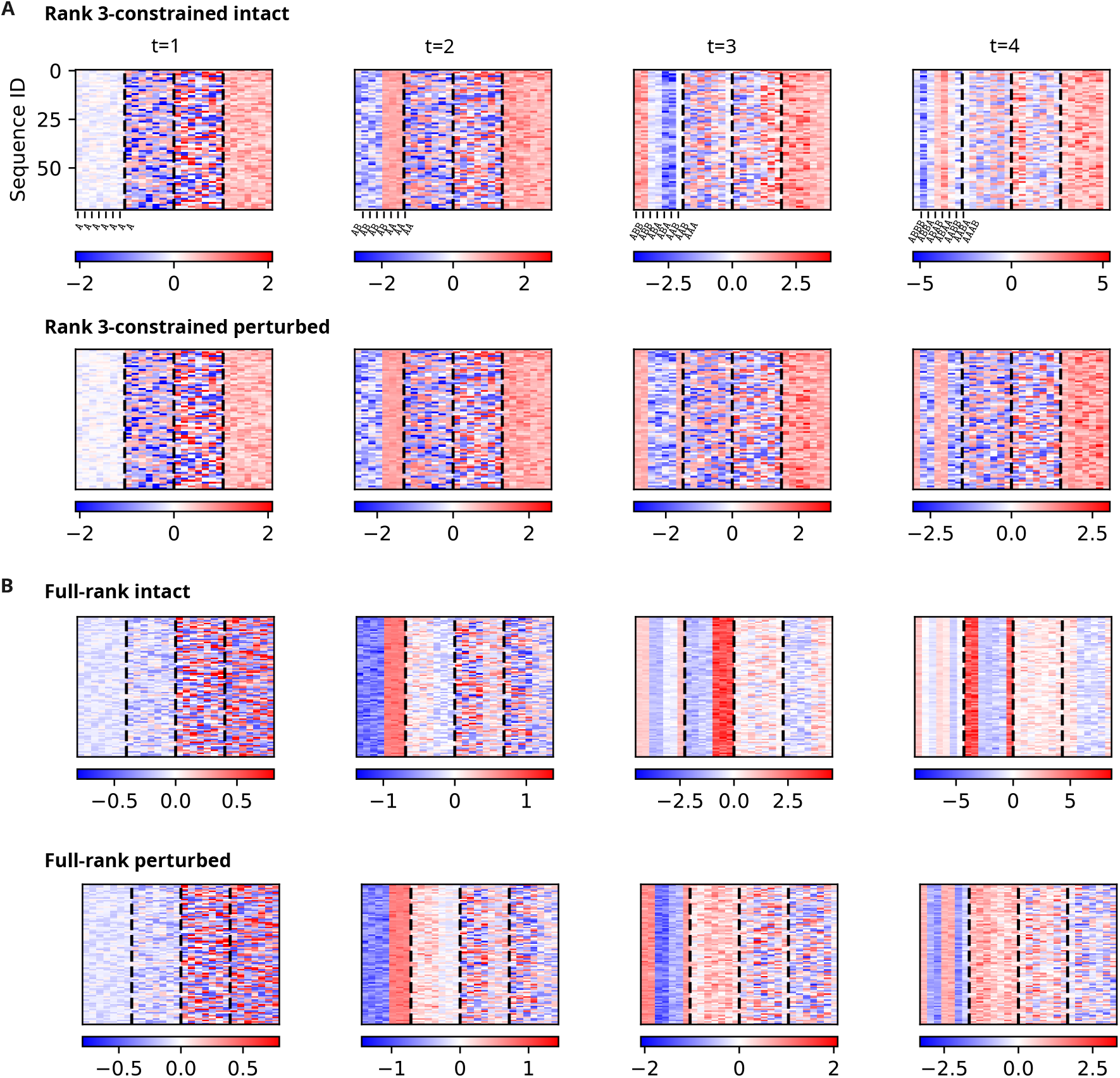
Perturbation experiments highlight the role of the singular components of the structured part of the recurrent connectivity. The RNN with a rank-3 recurrent weight matrix performs the task with high accuracy, approaching the performance of a full-rank one. In both cases, can see that the projections along the first principal components (**A**, top and **B**, top). By simulating a network where the dominant singular vector is removed from the recurrent weight matrix, we gain an insight into the role of the low-rank structured part (**A**, bottom and **B**, bottom). In both cases we notice the following. *i)* In the intact network the projections along the first singular components jointly carry information about the distinct classes into which the structure of sequences (or partial, processed so far) can be categorized. *ii)* When the dominant singular component is removed, the only projection that obviously carries information about the structure is SV1 itself: in particular, it is only informative of the latest seen transition (“same” vs “different”); since SV1 is removed from the recurrent weights, this is not propagated to the next time step, preventing the network from memorizing the whole structure.

**Figure 10:**
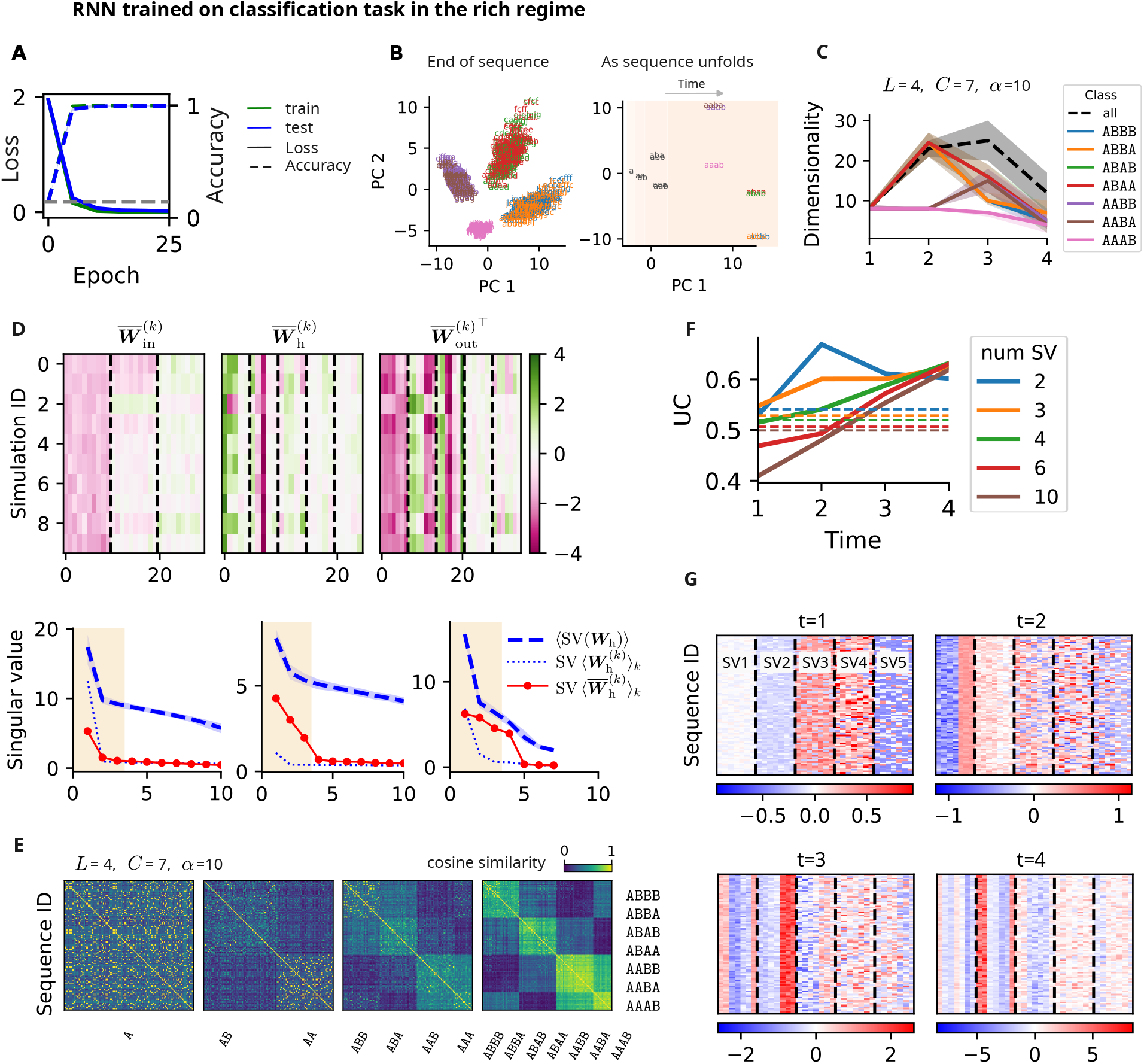
The results with “lazy” initialization regime hold in the “rich” case as well.

### 5.2 Transfer experiments

We studied three tasks using an RNN: one trained to classify sequences based on the abstract temporal structure of said sequences, a second trained to predict the next item in a sequence, and a third trained to reconstruct sequences.

#### Transferring classification pretrained weights to prediction network

While both classification and prediction trained networks exhibit good memory capacity as shown by their performance on sequences used in the training set, only the classification-trained network generalizes well across a wide range of parameters, particularly for longer sequences (Figs. 6 and 7). We found that the classification network achieves this via a progressive dimensionality reduction in its dynamics as the sequence unfolds (Fig. 2A, main text), which we linked to a low-rank structure in the recurrent connectivity matrix. In contrast, despite relying on similar sequence statistics (i.e. temporal patterns present in the classes), the prediction-trained network does not exhibit low-rank recurrent structure (Fig. 5D, main text). We note that one-step transition probabilities are not enough to perform the prediction task, and at any given time, the network requires knowledge of both the class and letter to perform this task. We hypothesized that the abstract structure learned by the classification network could be leveraged to improve prediction. To test this, we implemented multiple transfer learning protocols: either 1) initializing the prediction network’s weights and biases from a trained classifier and then freezing them (i.e. no more learning), or 2) initializing from the classifier and training afterwards. We applied these protocols separately to the input weights, recurrent weights, or both (Figs. 8A). When all parameters were frozen (protocol 1, red curve), prediction performance remained at chance (Fig. 8A). Freezing only the recurrent weights (blue curve) allowed for learning of memorization but not generalization. Freezing only input weights (pink) permitted learning but generalization was slow relative to learning from scratch (green).

Interestingly, when recurrent and input weights were transferred separately following protocol 2 (cyan and purple), we observed modest improvements in both memorization and generalization (Fig. 8A). However, combining both (input + recurrent; orange) led to sizeable gains in learning speed and generalization. To evaluate the robustness of this transfer, we ran this transfer protocol across a wide range of parameters, recurrent layer size *N*, number of classes *C* and sequence length *L* (Figs. 8B and C). Comparing against networks trained from scratch, we found that the transfer protocol 2 (input + recurrent) improved performance (in both memorization and generalization) early (epoch 10) and late (epoch 100) in training (green areas) for large-enough networks, but with no to negative effects in the performance of very small networks (red areas).

#### Transferring reconstruction pretrained weights to prediction network

To control that any improvement in generalization performance when transferring classification-pretrained weights to the prediction network is due to the abstract scaffold and not due to a “warm start” hypothesis, we trained a an autoencoder model. We then transferred its encoder weights following exactly the same protocol as before. We observe in this case that although the network learns faster in almost all the protocols in which weights are transferred, generalization does not improve, as in the classification-pretrained case. We conclude from this that faster learning can be attributed to the warm start provided by pretraining the network, but for better generalization, only the abstract scaffold results in transfer.

### 5.3 Parameters

We report here all the parameters required to reproduce the results. Table 1 recapitulates the parameters shared across all experiments in the manuscript, while Tables 2, 3 and 4 are specific to figures.

**Table 1:** Table of parameters shared across all experiments.

| Description | Size |
| --- | --- |
| Network |  |
| Learning rate | 0.001 |
| Batch size | 1 |
| Initialization | $\mathcal{U}\left(-\frac{1}{\sqrt{N}}, \frac{1}{\sqrt{N}}\right)$ |
| Transfer function | ReLU |
| Loss function | Cross-Entropy |
| Data |  |
| Number of unique letters per sequence $m$ | 2 |
| Size of alphabet $\alpha$ | 10 |

**Table 2:**
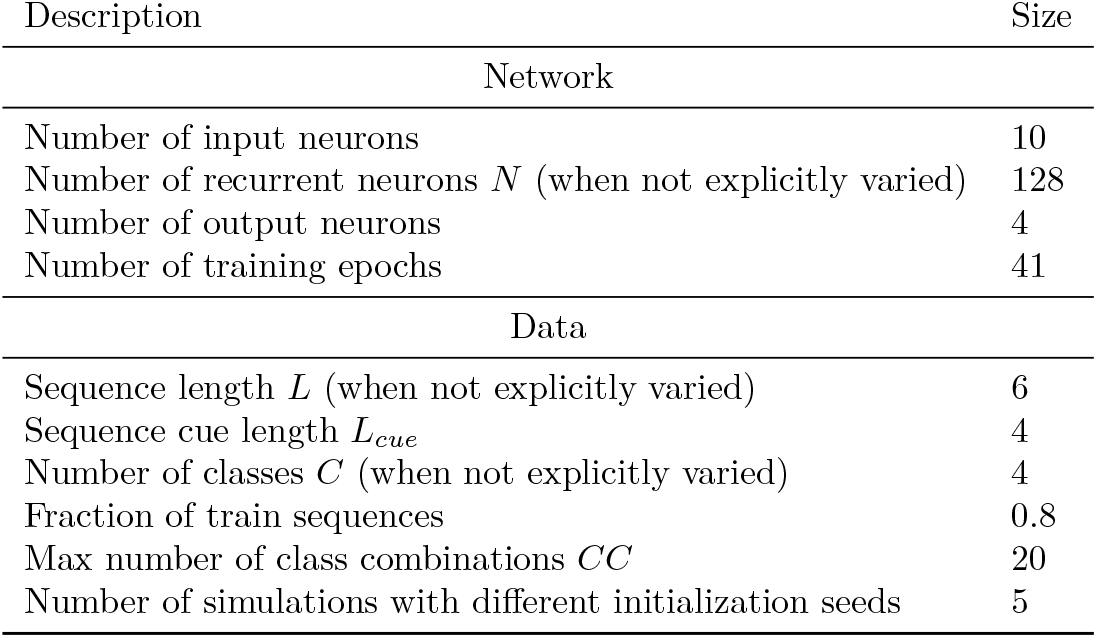
Table of parameters for Fig. 1 of main text.

**Table 3:**
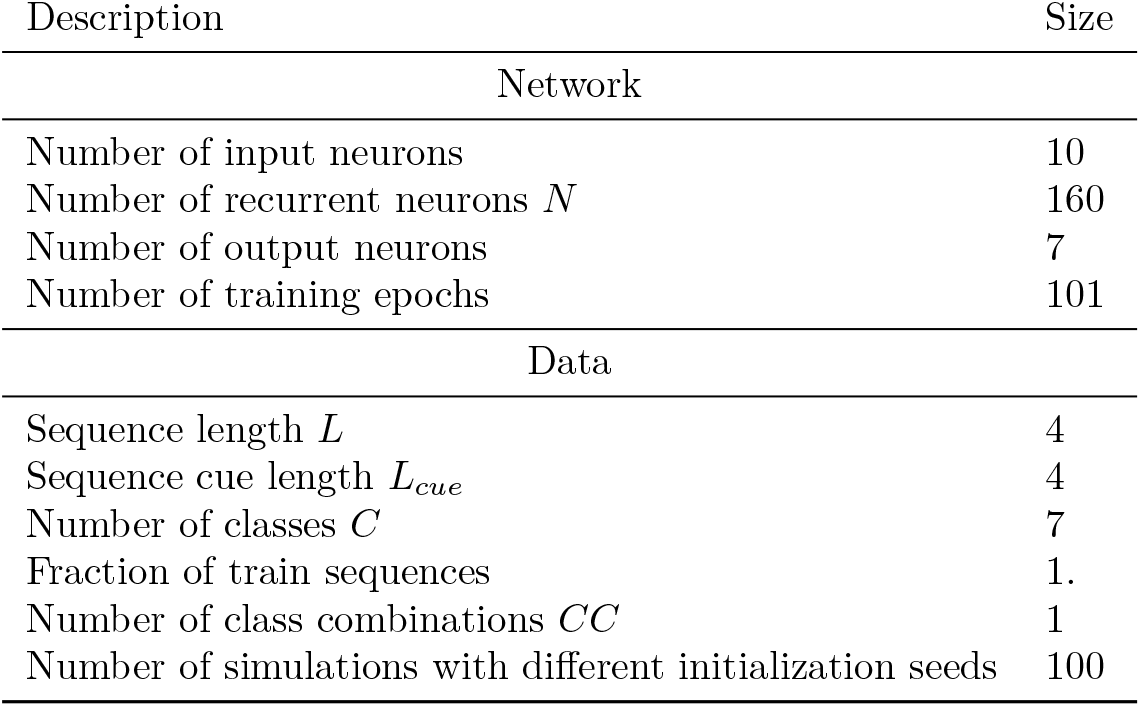
Table of parameters for Figs. 2 and 3 of main text.

**Table 4:** Table of parameters Fig. 4 of main text and Figs. 6, 7, and 8.

| Description | Size |
| --- | --- |
| Network |  |
| Number of input neurons | 10 |
| Number of recurrent neurons $N$ (when not explicitly varied) | 160 |
| Number of output neurons (class, pred) | 5, 10 |
| Number of training epochs (class, pred) | 41, 101 |
| Data |  |
| Sequence length $L$ (when not explicitly varied) | 9 |
| Sequence cue length $L_{cue}$ | 4 |
| Number of classes $C$ (when not explicitly varied) | 5 |
| Fraction of train sequences | 0.8 |
| Max number of class combinations $CC$ | 20 |
| Number of simulations with different initialization seeds | 5 |

## References

[1] Stanislas Dehaene, Florent Meyniel, Catherine Wacongne, Liping Wang, and Christophe Pallier. The neural representation of sequences: from transition probabilities to algebraic patterns and linguistic trees. Neuron, 88(1):2–19, 2015.

[2] Asaf Gilboa and Hannah Marlatte. Neurobiology of schemas and schema-mediated memory. Trends in cognitive sciences, 21(8):618–631, 2017.

[3] Frederic Charles Bartlett. Remembering: A study in experimental and social psychology. Cambridge university press, 1995.

[4] Adedamola Onih, Xinran Shen, Lida Pentousi, Vezha Boboeva, and Athena Akrami. The hippocampus enables abstract structure learning without reward. bioRxiv, pages 2026–02, 2026.

[5] Howard Eichenbaum. Hippocampus: cognitive processes and neural representations that underlie declarative memory. Neuron, 44(1):109–120, 2004.

[6] James CR Whittington, William Dorrell, Timothy EJ Behrens, Surya Ganguli, and Mohamady El-Gaby. A tale of two algorithms: Structured slots explain prefrontal sequence memory and are unified with hippocampal cognitive maps. Neuron, 113(2):321–333, 2025.

[7] Earl K Miller and Jonathan D Cohen. An integrative theory of prefrontal cortex function. Annual review of neuroscience, 24(1):167–202, 2001.

[8] Valerio Mante, David Sussillo, Krishna V Shenoy, and William T Newsome. Context-dependent computation by recurrent dynamics in prefrontal cortex. nature, 503(7474):78–84, 2013.

[9] Dorothy Tse, Rosamund F Langston, Masaki Kakeyama, Ingrid Bethus, Patrick A Spooner, Emma R Wood, Menno P Witter, and Richard GM Morris. Schemas and memory consolidation. Science, 316(5821):76–82, 2007.

[10] Dorothy Tse, Tomonori Takeuchi, Masaki Kakeyama, Yasushi Kajii, Hiroyuki Okuno, Chiharu Tohyama, Haruhiko Bito, and Richard GM Morris. Schema-dependent gene activation and memory encoding in neocortex. Science, 333(6044):891–895, 2011.

[11] Pierre Baraduc, J-R Duhamel, and Sylvia Wirth. Schema cells in the macaque hippocampus. Science, 363(6427):635–639, 2019.

[12] John Duncan. An adaptive coding model of neural function in prefrontal cortex. Nature reviews neuroscience, 2(11):820–829, 2001.

[13] A Schapiro and Nicholas Turk-Browne. Statistical learning. Brain mapping, 3(1):501–506, 2015.

[14] Gary F Marcus. The algebraic mind: Integrating connectionism and cognitive science. MIT press, 2003.

[15] Liping Wang, Lynn Uhrig, Bechir Jarraya, and Stanislas Dehaene. Representation of numerical and sequential patterns in macaque and human brains. Current Biology, 25(15):1966–1974, 2015.

[16] Shuchen Wu, Mirko Thalmann, and Eric Schulz. Two types of motifs enhance human recall and generalization of long sequences. Communications Psychology, 3(1):3, 2025.

[17] Robin A Murphy, Esther Mondragón, and Victoria A Murphy. Rule learning by rats. Science, 319(5871):1849– 1851, 2008.

[18] Gonzalo P Urcelay and Ralph R Miller. On the generality and limits of abstraction in rats and humans. Animal Cognition, 13:21–32, 2010.

[19] Chiara Santolin, Orsola Rosa-Salva, Lucia Regolin, and Giorgio Vallortigara. Generalization of visual regularities in newly hatched chicks (gallus gallus). Animal Cognition, 19:1007–1017, 2016.

[20] Michelle J Spierings and Carel Ten Cate. Budgerigars and zebra finches differ in how they generalize in an artificial grammar learning experiment. Proceedings of the National Academy of Sciences, 113(27):E3977–E3984, 2016.

[21] Sandra Mikolasch, Kurt Kotrschal, and Christian Schloegl. Transitive inference in jackdaws (corvus monedula). Behavioural processes, 92:113–117, 2013.

[22] Chiara Santolin and Jenny R Saffran. Constraints on statistical learning across species. Trends in Cognitive Sciences, 22(1):52–63, 2018.

[23] John P Cunningham and Byron M Yu. Dimensionality reduction for large-scale neural recordings. Nature neuroscience, 17(11):1500–1509, 2014.

[24] Peiran Gao and Surya Ganguli. On simplicity and complexity in the brave new world of large-scale neuroscience. Current opinion in neurobiology, 32:148–155, 2015.

[25] Mattia Rigotti, Omri Barak, Melissa R Warden, Xiao-Jing Wang, Nathaniel D Daw, Earl K Miller, and Stefano Fusi. The importance of mixed selectivity in complex cognitive tasks. Nature, 497(7451):585–590, 2013.

[26] Timothy EJ Behrens, Timothy H Muller, James CR Whittington, Shirley Mark, Alon B Baram, Kimberly L Stachenfeld, and Zeb Kurth-Nelson. What is a cognitive map? organizing knowledge for flexible behavior. Neuron, 100(2):490–509, 2018.

[27] Francesca Mastrogiuseppe and Srdjan Ostojic. Linking connectivity, dynamics, and computations in low-rank recurrent neural networks. Neuron, 99(3):609–623, 2018.

[28] Friedrich Schuessler, Alexis Dubreuil, Francesca Mastrogiuseppe, Srdjan Ostojic, and Omri Barak. Dynamics of random recurrent networks with correlated low-rank structure. Physical Review Research, 2(1):013111, 2020.

[29] Christian K Machens, Ranulfo Romo, and Carlos D Brody. Flexible control of mutual inhibition: a neural model of two-interval discrimination. Science, 307(5712):1121–1124, 2005.

[30] Omri Barak. Recurrent neural networks as versatile tools of neuroscience research. Current Opinion in Neurobiology, 46:1–6, 2017.

[31] Guangyu Robert Yang and Manuel Molano-Mazón. Towards the next generation of recurrent network models for cognitive neuroscience. Current Opinion in Neurobiology, 70:182–192, 2021.

[32] Laura Driscoll, Krishna Shenoy, and David Sussillo. Flexible multitask computation in recurrent networks utilizes shared dynamical motifs. bioRxiv, pages 2022–08, 2022.

[33] Matthew Botvinick and Takamitsu Watanabe. From numerosity to ordinal rank: a gain-field model of serial order representation in cortical working memory. Journal of Neuroscience, 27(32):8636–8642, 2007.

[34] Zhikun Chu, Qinghai Guo, Jie Cheng, Bo Ho, Si Wu, Yuanyuan Mi, et al. Learning and processing the ordinal information of temporal sequences in recurrent neural circuits. Advances in Neural Information Processing Systems, 36:33999–34020, 2023.

[35] Shuchen Wu, Stephan Alaniz, Shyamgopal Karthik, Peter Dayan, Eric Schulz, and Zeynep Akata. Concept-guided interpretability via neural chunking. arXiv preprint 2505.11576, 2025.

[36] Howard Eichenbaum. Time cells in the hippocampus: a new dimension for mapping memories. Nature Reviews Neuroscience, 15(11):732–744, 2014.

[37] Diederik P Kingma and Jimmy Ba. Adam: A method for stochastic optimization. arXiv preprint 1412.6980, 2014.

[38] Timo Flesch, Keno Juechems, Tsvetomira Dumbalska, Andrew Saxe, and Christopher Summerfield. Orthog-onal representations for robust context-dependent task performance in brains and neural networks. Neuron, 110(7):1258–1270, 2022.

[39] Alessandro Treves. On the perceptual structure of face space. BioSystems, 40(1-2):189–196, 1997.

[40] Vezha Boboeva, Romain Brasselet, and Alessandro Treves. The capacity for correlated semantic memories in the cortex. Entropy, 20(11):824, 2018.

[41] Elia Turner and Omri Barak. The simplicity bias in multi-task rnns: Shared attractors, reuse of dynamics, and geometric representation. Advances in Neural Information Processing Systems, 36, 2024.

[42] Niru Maheswaranathan, Alex Williams, Matthew Golub, Surya Ganguli, and David Sussillo. Universality and individuality in neural dynamics across large populations of recurrent networks. Advances in neural information processing systems, 32, 2019.

[43] Marlieke TR Van Kesteren, Dirk J Ruiter, Guillén Fernández, and Richard N Henson. How schema and novelty augment memory formation. Trends in neurosciences, 35(4):211–219, 2012.

[44] Marlieke TR van Kesteren, Mark Rijpkema, Dirk J Ruiter, and Guillén Fernández. Consolidation differentially modulates schema effects on memory for items and associations. PloS one, 8(2):e56155, 2013.

[45] Paul W Frankland and Bruno Bontempi. The organization of recent and remote memories. Nature reviews neuroscience, 6(2):119–130, 2005.

[46] Andrew M Saxe, James L McClelland, and Surya Ganguli. A mathematical theory of semantic development in deep neural networks. Proceedings of the National Academy of Sciences, 116(23):11537–11546, 2019.

